# Precision probabilistic mapping reveals personalized functional topography in the early postnatal human brain

**DOI:** 10.1101/2025.11.18.689151

**Authors:** Jianlong Zhao, Yuehua Xu, Zaixu Cui, Hongming Li, Lianglong Sun, Xinyuan Liang, Meizhen Han, Zilong Zeng, Qiongling Li, Tengda Zhao, Yong He

**Author notes:** Correspondence to: Tengda Zhao, Ph.D.,; Yong He, Ph.D.

## Abstract

Human adults exhibit remarkable individual differences in the brain’s functional topography, yet how such individualized topography emerges and shapes long-term cognition remains unknown. Using multimodal neuroimaging data of 419 infants (367 term-born; 52 preterm-born) from the Developing Human Connectome Project, we characterize personalized functional topographies in the neonate brain and evaluate their ability to predict 18-month neurocognitive outcomes. We find that typical functional topographies are present at birth, with interindividual variability organized along a sensorimotor-to-association hierarchy. These individualized maps encode early brain maturity and predict cognitive, language, and motor outcomes in early childhood, with the most predictive features localized to sensorimotor network boundaries. This functional refinement is structurally anchored by concurrent maturation in cortical myelination and sulcal depth. Preterm birth is associated with topographic alterations featuring atypical association network maturation. Our findings establish a foundational framework for precision modelling of early brain development and its influences on long-term cognitive trajectories.

## Introduction

The human brain exhibits substantial individual differences in the spatial architecture of its functional networks, or functional topography. Recent studies in adults ^1–8^ and youth ^9–11^ have suggested that these individualized topographies, represented as spatial loading maps of brain networks, are highly reliable and robust predictors of cognitive performance ^5, 9, 11^. This inter-individual variation follows a hierarchical sensorimotor-to-association gradient ^1, 4, 9, 12^ and correlates with fundamental cortical properties, including myelination patterns ^9^, gene expression profiles ^13, 14^, and evolutionary expansion ^9^. Despite these advances, the developmental origins of individualized functional topographies remain largely unknown. Addressing this gap is crucial to understand how functional architectures emerge early in life and shape subsequent neurocognitive and behavioural outcomes.

The early postnatal period is a critical phase of human brain development, characterized by rapid structural and functional reorganization ^15–17^. During this period, the characteristic architecture of functional networks becomes established in primary visual and somatomotor cortices, whereas higher-order association networks remain comparatively immature ^18, 19^. Although previous studies have reported individual differences in functional topography in early development ^15, 16, 20–23^, they possess several important limitations. First, functional connectivity patterns vary across infants ^20–23^, but personalized topographies have not been fully established. Although two recent studies ^24, 25^ generated personalized functional networks in infants using a hard parcellation framework; this approach fails to capture the spatially complex and heterogeneous functional configurations within specific networks ^10, 26–28^, which can be better characterized by probabilistic network modelling ^9, 10^. Moreover, combining term-born babies and preterm-born infants ^21, 22, 24^ in studies of early functional topologies may obscure normative developmental patterns, given that preterm birth is known to alter early brain development and elevate neurodevelopmental risks ^29–31^. Second, structural features such as myelination undergo rapid postnatal changes ^16, 32, 33^. However, how the structural maturation supports the development of individualized functional topography remains unclear. Third, despite that early functional organization provides a foundational neural architecture for long-term cognitive and behavioural capacities ^15, 16, 34^, the influence of personalized functional topographies emerging early in life on subsequent behavioural phenotypes remains unexplored.

To address these important questions, we leveraged the largest multimodal neonatal neuroimaging and 18-month follow-up neurocognitive dataset to date (367 term-born and 52 preterm-born) from the Developing Human Connectome Project (dHCP) ^35^. Using spatially regularized nonnegative matrix factorization (NMF) ^9, 11, 36^, we systematically investigated personalized functional topography during the early postnatal period. The specific aims of this study were to (1) delineate probabilistic functional topographies of neonatal brain networks, (2) quantify their interindividual variability, (3) develop a multivariable framework that uses topographic features to encode early brain maturation and predict 18-month neurocognitive outcomes, (4) elucidate the structural basis of early functional topographic refinements and (5) identify preterm-specific topographic alterations. Collectively, our work establishes a blueprint of personalized functional topographies in the first weeks of life, delineating their relationship with structural maturation and revealing their significant influence on later behavioural traits and health outcomes.

## Results

### Data samples

Following stringent quality control procedures (Fig. S1), we assembled high-quality multimodal neuroimaging, neurocognitive, and behavioural data from 419 term-equivalent infants (including 52 born preterm) sourced from the dHCP ^35^. The dataset was stratified into three subsets on the basis of prespecified analytic objectives. (i) Subset 1 (52 term neonates; gestational age [GA] at birth, 37–42 weeks; postmenstrual age [PMA] at scan, 37–44 weeks) data were used to generate initial group-level functional networks, which served as unbiased spatial constraints for subsequent personalized topographic analyses (Fig. 1a). (ii) Subset 2 (315 term neonates; GA at birth, 37–42 weeks; PMA at scan, 37–44 weeks) data were used to perform person-specific functional topographic mapping, including the quantification of individual variations, predictive modelling of brain maturity and 18-month neurodevelopmental outcomes, and establishment of the structural basis of early functional development (Fig. 1a–d). (iii) Subset 3 (52 preterm infants; GA at birth, 23–36 weeks; PMA at scan, 37–44 weeks) data were used to assess alterations in functional topography in preterm infants (Fig. 1e). For comparative neurodevelopmental benchmarking, we included task-free fMRI data from 200 unrelated young adults (22–25 years old) from the Human Connectome Project (HCP) ^37^. Demographic characteristics and imaging protocols are detailed in Table 1.

**Fig. 1.**
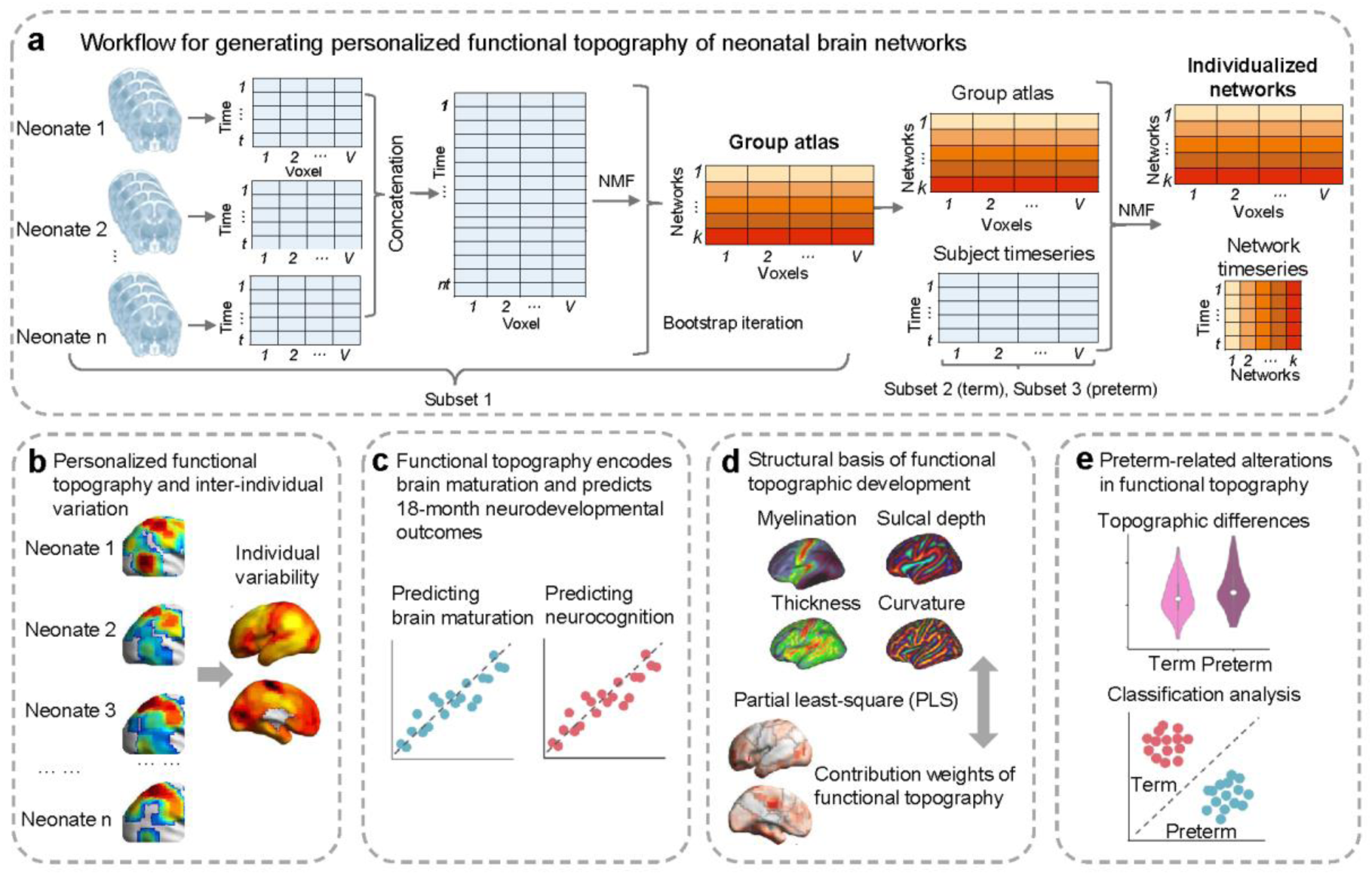
Study design and analysis flowchart. **a,** Workflow for generating group-based and personalized functional topographies of neonatal brain networks. **b,** Personalized functional topography and interindividual variation. **c,** Functional topography encodes brain maturation and predicts 18-month neurodevelopmental outcomes. **d,** Structural basis of functional topography by relating structural features (myelination, thickness, curvature, and sulcal depth) to the contribution weights of functional topography (brain-age and brain-behavioural prediction models) using partial least-square (PLS) analysis. **e,** Preterm functional topographic alterations in preterm infants were assessed through group comparisons and multivariable classification analyses.

**Table 1.** Infant demographics and follow-up neurodevelopmental data.

| **Group** | **Term** |  | **Preterm** |
| --- | --- | --- | --- |
| Subset | Subset 1 | Subset 2 | Subset 3 |
| Number of participants | 52 | 315 | 52 |
| Sex, num of females (%) | 36 (69.23%) | 134 (42.54%) | 19 (36.54%) |
| GA at birth (weeks) | 37-42 | 37-42 | 23-36 |
| PMA at scan (weeks) | 37-44 | 37-44 | 37-44 |
| Birth weight (kg) | 2.16-4.34 | 1.82-4.80 | 0.59-3.16 |
| Time points | 2300 | 2300 | 2300 |
| Scan time (mins) | 15 | 15 | 15 |
| **Follow-up** |  |  |  |
| Bayley III<sup>a</sup> - cognitive (S.D.) | N/A | 101.04 (11.16) | 101.49 (12.41) |
| Bayley III<sup>a</sup> - language (S.D.) | N/A | 98.37 (15.60) | 99.89 (17.28) |
| Bayley III<sup>a</sup> - motor (S.D.) | N/A | 102.50 (9.96) | 100.84 (9.71) |
| Q-CHAT<sup>b</sup> - (S.D.) | N/A | 30.33 (8.74) |  |

GA, gestational age; PMA, postmenstrual age; <sup>a</sup>Bayley Scales of Infant Development: Third Edition (Bayley-III) - # of complete assessments: 249 term-born neonates; <sup>b</sup>Quantitative Checklist for Autism in Toddlers (Q-CHAT) - # of complete assessments: 245 term-born neonates. Notably, Subset 1 was derived from dHCP Release 2, whereas Subsets 2 and 3 were selected from dHCP Release 3.

### Functional topography and individual variation in neonatal brain networks

To delineate person-specific functional networks in 315 term-born neonates (Subset 2), we employed NMF ^9, 11, 36^, a data-driven method that decomposes functional connectivity data into interpretable spatial components. Unlike hard partition approaches that assign each voxel exclusively to a single network, NMF generates probabilistic loadings for each voxel across networks (soft partition), reflecting graded contributions to distinct functional networks.

We first established a group-level functional atlas based on NMF using high-quality data from 52 term-born neonates (Subset 1; Fig. 1a). This group atlas served as an initial reference for individual-level decomposition, ensuring spatial correspondence across participants. Given the known differences between neonatal and adult functional architectures ^19^, we avoided the canonical 17-network adult atlas ^38, 39^ and instead adopted an 11-network neonatal resolution ^30^ (validated against alternative 9-/17-network resolutions in Fig. S2–S3). Our NMF analysis revealed well-defined neonatal sensorimotor networks (Fig. 2a), including a visual network (primary and lateral visual network: Networks 1–2) and a somatomotor network (foot motor network: Network 3; and hand and mouth motor network: Network 4). In contrast, high-order association networks exhibited marked immaturity. The default-mode network fragmented into anterior (DMN-a: Network 5), posterior (DMN-p: Network 6) and lateral (DMN-l and DMN-r: Networks 7-8) components. Frontoparietal and attentional networks appeared as isolated patches (e.g., superior parietal, dorsal frontal, and fronto-limbic: Networks 9-11). These probabilistic functional topographies in neonates align with the findings of prior reports describing well-developed sensorimotor networks and immature association networks during early postnatal development ^30, 40–43^; the present work extends these findings by establishing a probabilistic parcellation framework for neonatal brains. For visualization, we generated a group-based parcellation atlas by assigning each voxel to the network with the highest loading in the probabilistic maps (Fig. 2b). Using the group atlas as a spatial constraint, we implemented individualized NMF to derive eleven person-specific functional networks for 315 neonates in Subset 2 (Subset 1 was excluded to avoid data leakage). To validate the efficacy of individualized network parcellations, we assessed the functional homogeneity of each network, quantified as the average Pearson’s correlation coefficients of time series between all pairs of voxels within a given network. Compared with the group-level atlas, individualized networks exhibited greater functional homogeneity (*t*(314) = 20.70, *P* = 2.33 × 10⁻⁷, Cohen’s *d* = 1.98, 95% CI: [0.03, 0.06]; Fig. 2c).

**Fig. 2.**
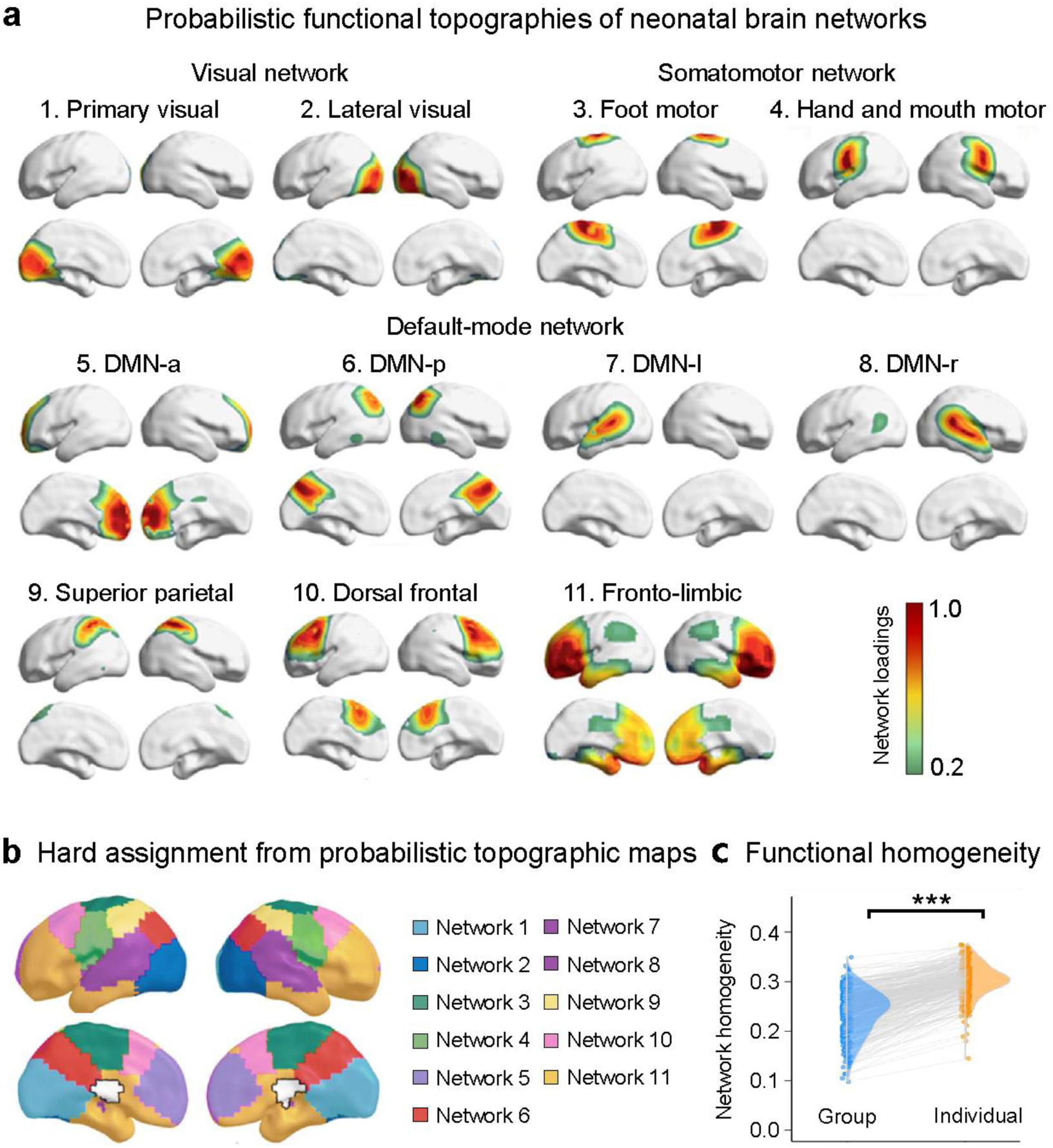
Functional topographies of neonatal brain networks. **a**, Group-level probabilistic network topographies. Voxel-wise loadings represent the degree of network membership, with warmer colours indicating higher loadings. **b,** Hard parcellation obtained by assigning each voxel to the network with the highest loading. **c,** Compared with the group-level atlas, the individualized network maps exhibited significantly higher within-network functional homogeneity (*t*(314) = 20.70, *P* = 2.33 × 10⁻⁷, Cohen’s *d* = 1.98, 95% CI: [0.03, 0.06], *n* = 315, two-sided paired t-test). ***, *P* < 0.001. DMN-p, posterior DMN; DMN-a, anterior DMN; DMN-l. left DMN; DMN-r, right DMN. Source data are provided as a Source Data file.

We next investigated interindividual variability in neonatal functional topography. Visual inspection revealed that individual variations were apparent in the association cortices, such as those in the dorsal frontal network (Network 10), and that the primary networks, such as the foot motor network (Network 3), were broadly conserved across neonates (Fig. 3a). To quantify these interindividual variabilities in functional topography, we computed the median absolute deviation (a nonparametric statistic of variance) of probabilistic loading across neonates for each network ^9, 44^. We found that the association cortices exhibited greater interindividual topographic variability than the primary cortices (*P*_spin_ = 0.017) (Fig. 3b). Further analyses of four topographical properties—position, size, overlap, and homogeneity ^5, 45^—also confirmed that the association networks were more variable across neonates than the primary networks (position: *P*_spin_ = 0.002; size: *P*_spin_ = 0.026; overlap: *P*_spin_ = 0.0004; homogeneity: *P*_spin_ = 0.0006; Fig. 3c and Fig. S4). Notably, this neonatal variability pattern was spatially correlated with adult interindividual differences (HCP dataset, r = 0.46, *P*_spin_ < 0.001; Fig. 3d), suggesting a conserved organizational hierarchy. Importantly, our findings trace the origins of these topographical variations, offering the earliest evidence for their ontogenetic establishment. Finally, to delineate fine-grained variability patterns along the cortical sheet, we performed boundary concentration analysis by assessing the spatial correlations between individual variability maps and network loading maps. We observed significant negative correlations at the global (r = −0.61, *P*_spin_ < 0.001) and network levels (all *P*_spin_ < 0.05, except Network 10) (Fig. 3e), indicating that the greatest interindividual variations tended to be located at network boundaries where the network loadings were smaller. This effect was more pronounced in the primary networks than in the association networks (*P*_spin_ = 0.031; Fig. 3e).

**Fig. 3.**
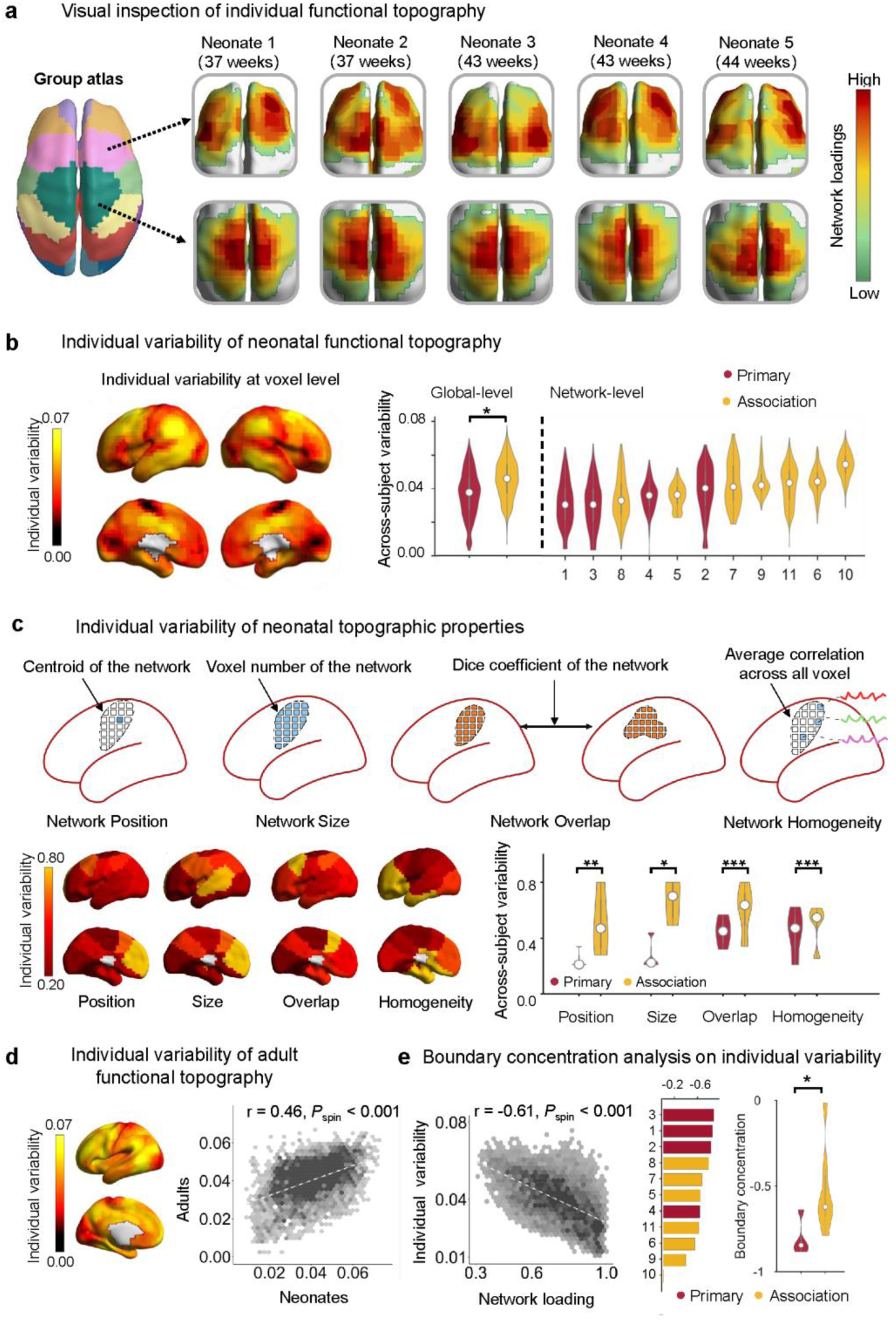
Interindividual variability in neonatal functional topography. **a,** Visual inspection of individualized voxel-wise probabilistic network maps in representative neonates. **b,** Quantification of interindividual variability of neonatal functional topography at the global and network levels. Association networks exhibited greater interindividual variability than the primary networks at the global level (*P*_spin_ = 0.017, one-sided). **c,** Quantification of interindividual variability of four topographical metrics, including position, size, overlap, and homogeneity. Association networks exhibited greater interindividual variability than primary networks across all four topographical properties (position: *P*_spin_ = 0.002; size: *P*_spin_ = 0.026; overlap: *P*_spin_ = 0.0004; homogeneity: *P*_spin_ = 0.0006, uncorrected p values, one-sided). **d,** Spatial correlation of functional topographic maps between neonatal and adult brains (*r* = 0.46, *P*_spin_ < 0.001, one-sided). **e,** Boundary concentration analysis by correlating interindividual topographic variability and network loadings of functional topography at both the global and network levels (*r* = 0.61, *P*_spin_ < 0.001, one-sided). For all violin plots, the central white point indicates the median, black lines represent the interquartile range (25th–75th percentiles), whiskers extend to the most extreme data points within 1.5 × IQR from the quartiles, asterisks indicate statistical significance (*, *P* < 0.05; **, *P* < 0.01; ***, *P* < 0.001). Source data are provided as a Source Data file.

### Functional topography undergoes refinement in the early postnatal period and encodes brain maturity

Having identified the interindividual topographic variability in neonatal functional networks, we next investigated how person-specific functional topography encodes brain maturation in 315 neonates (Subset 2).

First, we performed mass univariable analyses to characterize developmental changes in both total and voxel-wise network organization. (i) For the total network representation, we summed voxel-wise loadings for each probabilistic network and modelled age-related changes using generalized additive models (GAMs), controlling for sex, mean framewise displacement (FD), and scan-birth age intervals. The lateral visual network (Network 2) exhibited a nonlinear increase in total representation with age (*P* = 0.0003, *P*_FDR_ = 0.004), whereas the fronto-limbic network (Network 11) gradually decreased with age (*P* = 0.004, *P*_FDR_ = 0.02) (Fig. 4a). (ii) Voxel-wise GAM analyses for each probabilistic network revealed significant age-dependent refinement in network loadings in the visual (Networks 1–2), somatomotor (Network 4), superior parietal (Network 9), and frontal-limbic (Network 11) networks (Gaussian random field (GRF) correction at voxel-level *P* < 0.001, cluster-level *P* < 0.05) (Fig. 4b). No significant sex effects were observed for either global or voxel-wise functional topography, although at an uncorrected level, the total size of lateral visual network was larger in term-born males than females (z = 2.78, *P* = 0.005). Given the narrow age range of approximately 5 weeks across the neonatal cohort, we also employed a linear model to characterize age-related changes in network loading, which yielded results compatible with those from the GAM analyses (Fig. S5).

**Fig. 4.**
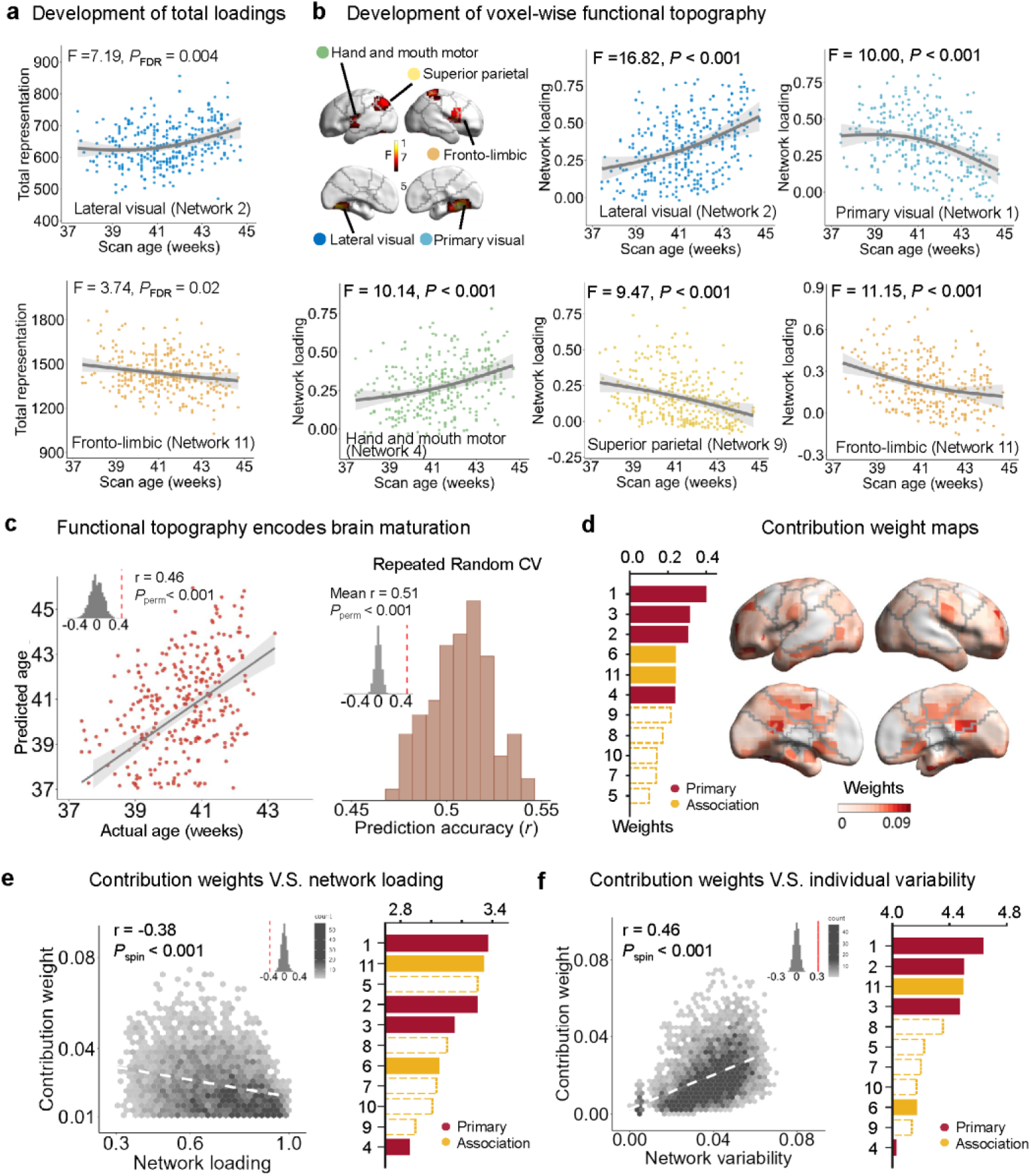
Functional topography undergoes progressive refinement and encodes brain maturity during the early postnatal period. **a,** Developmental trajectories of total network representation across scan ages (*n* = 315 term-born neonates, FDR corrected q < 0.05, corresponding to uncorrected *P* = 0.004, two sided). The total representation was quantified as the sum of the voxel-wise probabilistic loadings for each network. The scatter plot shows the fitted developmental curves with the shaded area indicating the 95% confidence interval around the fitted mean. **b,** Developmental trajectories of the voxel-wise functional topography. Analyses were conducted using two-sided tests at the voxel level (uncorrected *P* < 0.001) and corrected at the cluster level (GRF correction at *q* < 0.05) using Gaussian random field theory. The scatter plot shows the fitted developmental curves with the shaded area indicating the 95% confidence interval around the fitted mean. **c,** Functional topography encodes brain maturation using Ti-PCA and SVR frameworks. The scatter plot shows the prediction performance (r = 0.46, *P*_spin_ < 0.001, one-sided). **d,** Network- and voxel-level contribution weights derived from the brain-age prediction model. **e,** Spatial correlations between contribution weight and network loading across voxels (*r* = 0.38, *P*_spin_ < 0.001, one-sided). **f,** Spatial correlation between contribution weight and interindividual topographic variability (*r* = 0.46, *P*_spin_ < 0.001, one-sided). For all scatter plots, the fitted curves with the shaded area indicate the 95% confidence interval around the fitted mean. Primary and association networks are colour coded. Dashed outlines denote networks that were not classified as high-contributing networks (z-score < 0). Source data are provided as a Source Data file.

Second, to capture the high-dimensional reconfiguration of the functional topography, we developed a multivariable prediction framework that combines topography-independent principal component analysis (Ti-PCA) with support vector regression (SVR) (Fig. S6). Briefly, PCA was performed separately within each network in the training set while preserving feature independence across networks. The resulting PCA components were integrated into an SVR model with linear kernel as feature sets to predict postmenstrual age at scan. Using 10-fold cross-validation, we evaluated prediction accuracy via the Pearson’s correlation coefficient between predicted age (“brain maturity index”) and chronological age in the testing set after regressing out the effects of sex, mean FD, and scan-birth age intervals. This model achieved significant prediction accuracy (r = 0.46, permutation test, *P*_perm_ < 0.001). Robustness was confirmed across 100 repeated random folding analyses, yielding an averaged accuracy of r = 0.51 and mean absolute error (MAE) = 0.84 (*P*_perm_ < 0.001, Fig. 4c). Although single-network features also provided significant predictions (r = [0.15–0.40], *P*_perm_ < 0.002; Fig. S7), their inferior performance relative to that of TiPCA underscores the value of multinetwork integration for modelling brain maturity. High-contributing networks—identified by combining feature weights and cross-fold stability (z-scores > 0)—included all primary sensorimotor networks (Networks 1–4) and specific association networks (e.g., DMN-p, Network 6, and the fronto-limbic network, Network 11) (Fig. 4d). Voxel-wise feature weights were negatively correlated with network loading (r = −0.38, *P*_spin_ < 0.001; Fig. 4e) and positively correlated with interindividual variability (r = 0.46, *P*_spin_ < 0.001; Fig. 4f). Taken together, these results suggest that early postnatal development is primarily encoded at the network boundaries of the sensorimotor system and a few association systems and is further shaped by person-specific network variability.

### Neonatal functional topography predicts neurodevelopmental outcomes at 18 months

We next assessed whether personalized functional network topography at birth could predict individual neurodevelopmental outcomes at 18 months. In Subset 2, neonates (n = 249) participated in a follow-up visit at 18 months, where cognitive, language, and motor abilities were evaluated using the Bayley Scales of Infant and Toddler Development, 3rd Edition (Bayley-III) ^46^. Univariable GAM analyses revealed associations between total network loading and 18-month neurodevelopmental outcomes (Networks 4, 5, and 8, uncorrected p = 0.015∼0.048, covariates: sex, mean FD, scan age, and scan-birth age interval, Fig. S8), though these did not survive FDR correction. Multivariable analysis using Ti-PCA and SVR (linear kernel) revealed that the individual functional topography predicted individual cognitive (r = 0.39, *P*_perm_ < 0.001), language (r = 0.29, *P*_perm_ < 0.001), and motor (r = 0.37, *P*_perm_ < 0.001) scores (10-fold cross-validation; covariates: sex, mean FD, scan age, and scan-birth age intervals, at FDR corrected q < 0.001) (Fig. 5a). This prediction remained robust following 100 random-fold validations (cognition: mean r = 0.34; language: mean r = 0.27; motor: mean r = 0.34; all *P*_perm_ < 0.001; Fig. 5a) and outperformed single-network predictions (cognition: r range = [0.10, 0.32]; language: r range = [0.11, 0.27]; motor: r range = [0.13, 0.31]; Fig. S9-11). For neonates (n = 245) in Subset 2, the Quantitative Checklist for Autism in Toddlers (Q-CHAT) was completed at the 18-month follow-up visit to assess atypical behavioural risk. Multivariable analysis using Ti-PCA and SVR (linear kernel) revealed that person-specific functional topography predicted individual Q-CHAT scores (10-fold cross-validation: r = 0.29, *P*_perm_ < 0.001; 100 random-fold validation: mean r = 0.30, *P*_perm_ < 0.001, at FDR corrected q < 0.001) (Fig. 5a).

**Fig. 5.**
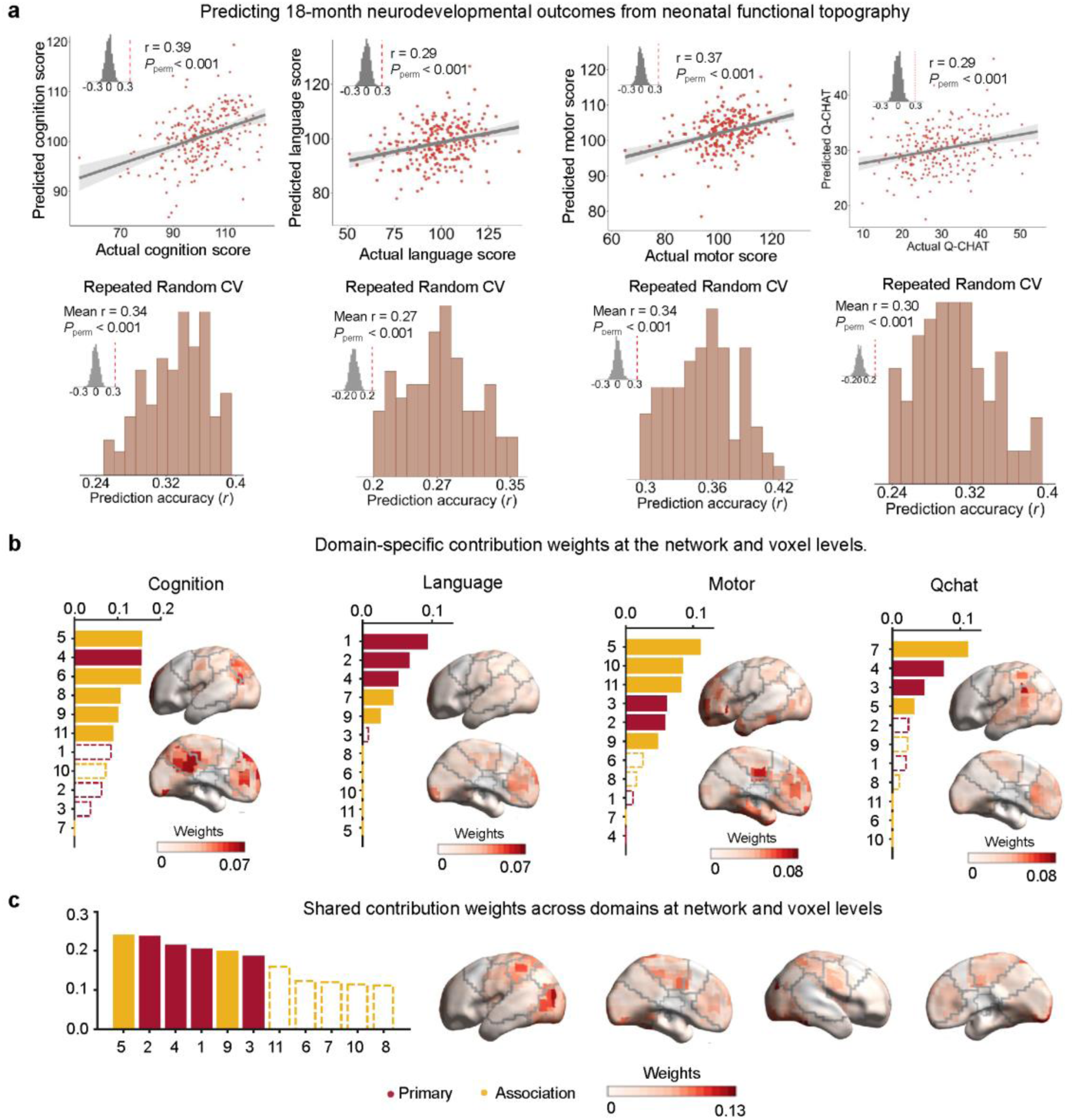
Neonatal functional topography predicts neurodevelopmental outcomes at 18 months. **a,** Multivariable prediction of neurodevelopmental outcomes at 18 months using personalized neonatal functional topography (Bayley-III: *n* = 249 term-born neonates; Q-CHAT: *n* = 245 term-born neonates). The scatter plot shows the prediction performance (all *P*_spin_ < 0.001, all FDR corrected *q* < 0.001, one-sided). Fitting curves with the shaded area indicate the 95% confidence interval around the fitted mean. The bottom panels display the prediction accuracy distributions across 100 repeated random cross-validation iterations. **b,** Domain-specific contribution weights at the network and voxel levels. The bars indicate network-level weights (red: primary; yellow: association). Dashed outlines denote networks that were not classified as high-contributing networks (z-score < 0). **c,** Shared contribution weights across domains at network and voxel levels. The bars indicate network-level weights (red: primary; yellow: association). Dashed outlines denote networks that were not classified as high-contributing networks (z-score < 0). Source data are provided as a Source Data file.

Next, we delineated domain-specific and domain-shared high-contributing voxels across all neurodevelopmental and behaviour scores. After regressing out the contribution maps of other domains, we observed distinct spatial patterns for each domain (Fig. 5b): cognition-specific contributions were primarily concentrated in several association networks (e.g., DMN-a, Network 5, and DMN-p, Network 6) and hand and mouth network (Network 4); language-specific contributions localized primarily to visual (Networks 1 and 2) and hand and mouth regions (Network 4); motor-specific contributions mapped mainly to the DMN-a (Network 5), dorsal frontal (Network 10), and somatomotor network (Networks 2 and 3); and Q-CHAT-related contributions were predominantly found in the DMN-l (Network 7) and somatomotor regions (Networks 3 and 4) (Fig. 5b). Voxel-wise contributing weights were negatively correlated with network loading (cognition: r = −0.39; language: r = −0.40; motor: r = −0.46; Q-chat: r = −0.32; all *P*_spin_ < 0.001) and positively correlated with individual variability (cognition: r = 0.48; language: r = 0.48; motor: r = 0.52; Q-chat: r = 0.39; all *P*_spin_ < 0.001) (Fig. S12). Finally, applying principal component analysis (PCA) to all contribution maps revealed a shared subset of networks that consistently contributed across domains, including the visual (Networks 1 and 2), somatomotor (Networks 3 and 4), DMN-a (Network 5), and superior parietal (Network 9) regions (Fig. 5c). Voxel-wise feature weights shared across domains were negatively correlated with interindividual variability (r = −0.46, *P*_spin_ < 0.001). Taken together, these results suggest that the functional topography-based predictions for neurodevelopmental outcomes were driven mainly by network boundaries with high across-neonate variability.

### Functional topographic refinement in the neonatal brain is anatomically constrained by structural maturation

We sought to delineate whether neonatal functional topographic refinement during the early postnatal period is anatomically constrained by cortical structural properties. Using structural images from 301 term-born neonates (Subset 2), we first generated individual cortical maps of four anatomical features, namely, cortical myelination, cortical thickness, surface curvature, and sulcal depth. We subsequently used GAMs to examine the spatial maturation of these structural features during the early postnatal period, with scan age as the main factor and sex, mean FD, and scan-birth age interval as covariates (Fig 6a). Cortical myelination exhibited pronounced age-dependent changes, particularly in the sensorimotor and insular cortex (*P* = 0.02, *P*_FDR_ < 0.05). Cortical thickness, curvature, and sulcal depth exhibited age-related changes in the boundaries of the medial foot motor cortex and anterior DMN (*P* = 0.015, *P*_FDR_ < 0.05).

**Fig. 6.**
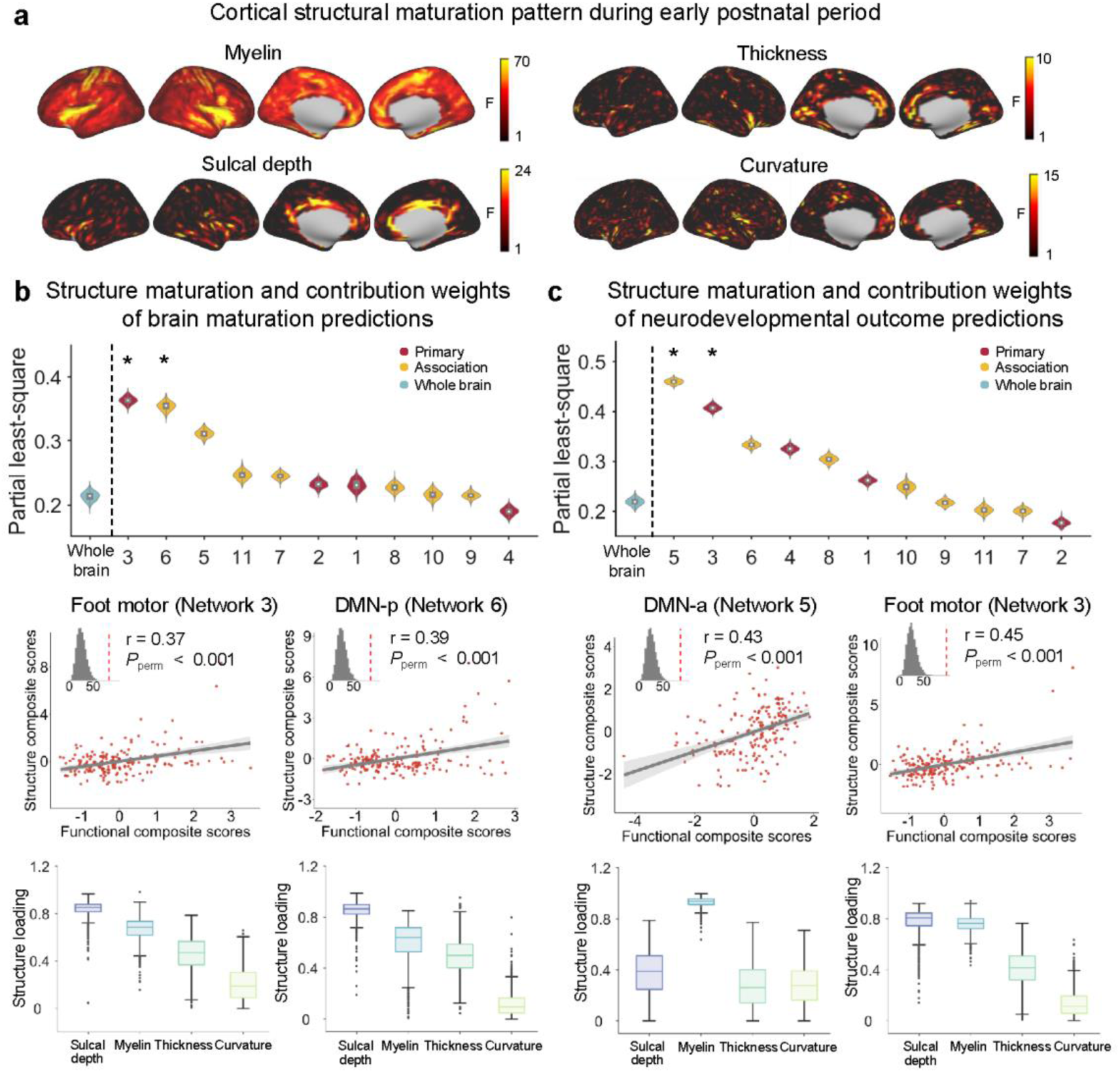
Functional topographic refinement in the neonatal brain is structurally constrained by cortical anatomical maturation. **a,** Cortical structural maturation pattern during the early postnatal period (*n* = 301 term-born neonates). F statistics for the effects of scan age are shown for myelination, sulcal depth, cortical thickness, and curvature (uncorrected *P*). **b,** Associations between structural maturation and functionally defined, network-level contribution weights from the brain-age prediction models. **c,** Associations between structural maturation and contribution weights from the brain–behaviour prediction models. In panels (b) and (c), violin plots display whole-brain and network-wide structure‒function associations across 1000 bootstrap resamples (asterisks represent the significance at FDR corrected *q* < 0.01, corresponding to uncorrected *P* = 0.0001, one sided). Red and yellow violins represent primary and association networks, respectively. Scatter plots show structure–function correlations at the network level (the fitting curves with the shaded area indicate the 95% confidence interval around the fitted mean, all *P*_perm_ < 0.001 at uncorrected level, one-sided). Box plots depict structural loadings for sulcal depth, myelination, thickness, and curvature across 1,000 bootstrap resamples. For all violin and box bars, the median is showed as the white circle (violins) or the center line (boxes), the lower and upper bounds of the bar correspond to the 25th and 75th percentiles, the whiskers extend to the most extreme data points within 1.5 × IQR. DMN-p, posterior DMN; DMN-a, anterior DMN. Source data are provided as a Source Data file.

Next, we performed partial least-square (PLS) analyses to assess the latent relationship between these maturation maps (*F* values of the age effect) and the contribution weights of network-specific functional topography in the brain-age and brain-behaviour prediction models. To mitigate potential biases due to the unequal network size, we applied bootstrap resampling (100 iterations) to ensure the same number of voxels across networks prior to the PLS analysis. The contributing feature weights of functional networks in brain age were correlated with age-related structural maturation (foot motor: r = 0.37, *P*_perm_ < 0.001; DMN-p: r = 0.39, *P*_perm_ < 0.001; *P*_FDR_ < 0.03), with the greatest loading value at the sulcal depth and in the myelin maps (Fig 6b). The contributing feature weights of functional networks in brain-behavioural predictions were correlated with structural maturation (DMN-a: r = 0.43, *P*_perm_ < 0.001; foot motor: r = 0.45, *P*_perm_ < 0.001; *P*_FDR_ < 0.03), with myelination and sulcal depth emerging as the strongest loading factors (Fig. 6c). These results suggest that the refinements of functional topography in the neonatal brain are structurally supported by cortical maturation, particularly by myelination and sulcal depth.

### Altered network topography and functional maturation in preterm infants

Previous studies have demonstrated that compared with term-born controls, preterm infants exhibit altered patterns of functional brain maturation ^30, 31, 47^. Here, we systematically investigated functional topographic alterations among 52 preterm infants scanned at term-equivalent age (Subset 3). For comparison, we used the neuroimaging data of 73 term-born neonates (Subset 2), matched for scan age, sex, and mean FD (Table S1). Preterm infants had lower birth weights than term-born controls did (z = 8.7; *P* = 3.22 × 10⁻^18^, Cohen’s d = 2.64) but had comparable 18-month neurodevelopmental outcomes (Table S1). The Ti-PCA-based support vector classification (linear kernel) analysis robustly distinguished preterm infants from term infants, achieving 82% accuracy (sensitivity: 75%; specificity: 86%; 1000 permutation tests: *P*_perm_ < 0.001; 100 random fold cross validations: mean accuracy = 79%, *P*_perm_ < 0.001) after controlling for sex, mean FD, and scan age (Fig. 7a). The most discriminative networks were primarily in several association systems, including the fronto-limbic network (Network 11), superior parietal network (Network 9), and DMN (Networks 5–7) (Fig. 7a). These voxel-wise contribution weights were negatively correlated with network loading (r = −0.36, *P*_spin_ < 0.001) and positively correlated with individual variability (r = 0.48, *P*_spin_ < 0.001) (Fig. S13c, d). Mass univariable GAM analyses (covariates: sex, mean FD, and scan age) also confirmed atypical functional topography in these association networks in preterm infants compared with term-born controls (Fig. S13a, b). We further assessed whether preterm infants exhibited functional topographic maturation distinct from that of term infants (Fig. 7b). Using the TiPCA model trained on the full sample of term-born neonates, we obtained significant predictions for preterm infants (r = 0.41, *P* = 0.002 for the whole brain; r = [0.31, 0.39], *P* <= 0.02 for specific networks) (Fig. S13e). After the personalized brain age gap (BAG, defined as predicted minus actual age) was adjusted by the slope and intercept of the predicted vs. actual age fitting^48^ (Fig. S14), preterm infants exhibited greater BAGs than term-born infants (whole-brain: t = 2.37, *P* = 0.018, Cohen’s *d* = 0.36; network-level: DMN-p: t = 3.71, *P* = 0.0002, Cohen’s d = 0.56; frontal-limbic: t = 3.39, *P* = 0.0008, Cohen’s d = 0.51) (Fig. 7b). These findings reflect altered topographic maturation in preterm infants during early postnatal development. The DMN-p BAG was positively correlated with 18-month motor scores in preterm infants (r = 0.54, *P* = 0.001, *P*_FDR_ = 0.02; covariates: sex, mean FD, scan age, and the scan-birth interval) (Fig. 7b), but the whole-brain BAG and other network BAG were not significantly correlated. Additionally, we applied the models trained on term-born infants to predict neurodevelopmental outcomes and Q-CHAT scores in the preterm cohort (Subset 3). However, none of these models yielded significant predictions (all *P*_perm_ > 0.11), suggesting potentially distinct mechanisms underlying neurodevelopment in preterm versus term infants.

**Fig. 7.**
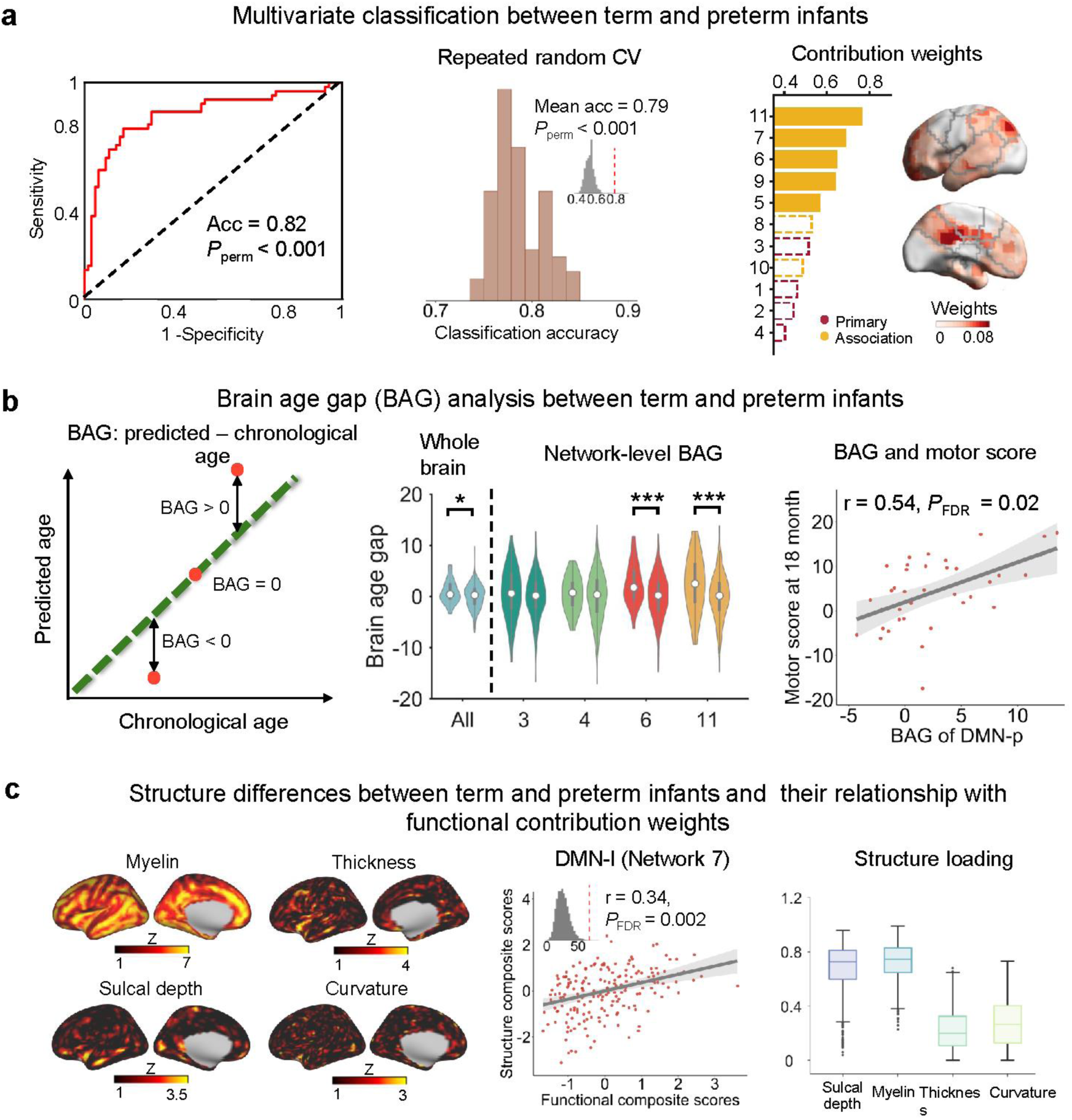
Altered network topography and functional maturation in preterm infants. **a,** Multivariable classification between term and preterm infants on the basis of individualized functional topography (Term infants: *n* = 73; Preterm infants: *n* = 52). Classification performance was evaluated using 10-fold cross-validation and repeated across 100 random folds (*P*_perm_ < 0.001, one-sided). Network- and voxel-level contribution weights are shown. **b,** Brain age gap analysis between term and preterm infants. Left: the BAG was computed as the predicted age minus chronological age. Middle: the difference of BAG between preterm (the left violin) and term (the right violin) infants (asterisks indicate statistical significance at *, *P* < 0.05; ***, *P* < 0.001, uncorrected paired t-test, two-sided). Right: association between the BAG and motor scores at 18 months (*r* = 0.54, FDR corrected *q* < 0.05, corresponding to uncorrected *P* = 0.0006, one-sided). The scatter plot shows the fitting curves with the shaded area indicating the 95% confidence interval around the fitted mean. **c,** Structural differences between term and preterm infants across four cortical features and their relationship with contribution weights of functional topography. Left: the distribution of group differences (Z scores) between term and preterm infants on myelination, sulcal depth, cortical thickness, and curvature. Middle: PLS analysis reveals the significant association between structural differences and functional contribution weights of the classification (DMN-l network, *r* = 0.34, FDR corrected *q* < 0.05, corresponding to uncorrected *P* = 0.0002, one-sided). The fitting curves with the shaded area indicate the 95% confidence interval around the fitted mean. Right: boxes depict structural loadings for sulcal depth, myelination, thickness, and curvature across 1,000 bootstrap resamples, the median shown as the center line, the lower and upper bounds of the boxes correspond to the 25th and 75th percentiles, the whiskers extend to the most extreme data points within 1.5 × IQR, and data points beyond the whiskers are shown as outliers. DMN-l, left DMN; BAG, brain age gap. Source data are provided as a Source Data file.

Finally, structural analyses using univariable GAMs revealed significantly lower myelination in the orbital and medial frontal and temporal parietal cortex in preterm infants than in term-born controls (*P* = 0.04, *P*_FDR_ < 0.05) (Fig. 7c). Cortical thickness, curvature, and sulcal depth exhibited age-related changes in the boundaries of the dorsal frontal and temporal cortex (*P* = 0.04, *P*_FDR_ < 0.05). Multivariable PLS analysis was used to test whether structural differences (*z* values) were correlated with the functionally defined feature contributions of the classification model. We showed that the DMN-l contribution weights were correlated with structural abnormalities (DMN-l: r = 0.34, *P*_FDR_ = 0.002), which were primarily driven by the cortical development of myelination and sulcal depth (Fig. 7c, S13f).

### Sensitivity analyses

Our results were validated using multiple analytical strategies, including assessments of sample size, scan duration, network resolution, and head motion control. (i) To evaluate the effects of head motion, the analysis was repeated using neonatal data under a stricter FD threshold (mean FD < 0.3 mm) (Fig. S15). (ii) To examine the influence of scan duration, we performed a non-overlapping split-half evaluation, in which functional networks reconstructed from one half of the total scan (both random and sequential samplings with progressively increasing durations) were compared against those derived from the remaining half ^38, 39^ (Fig. S16). (iii) To assess the impact of sample size on brain age and brain behaviour prediction models ^49, 50^, bootstrapped subsampling was performed across logarithmically spaced sample sizes (*n* = 25–284 neonates for brain age; *n* = 25–224 neonates for behaviour), with prediction analyses repeated 100 times per sample size (Fig. S17). (iv) To test the reproducibility of the results, a split-half validation approach was employed, in which subsets were matched by neonatal scan age (Fig. S18). (v) To determine the effect of sample selection on initial group-level atlases, we randomly selected subsets of term-born neonates (ranging from 10 to 300 scans) (Fig. S19). (vi) To evaluate sensitivity to network resolution, neonatal functional topographies were regenerated using parcellation schemes with k = 9 ^41, 51^ and k = 17 ^38, 52, 53^ (Fig. S20). (vii) To evaluate the covariate regression strategy, we repeated the prediction analysis by estimating covariate effects exclusively within the training folds rather than across the full sample (Fig. S21). (viii) To mitigate potential biases arising from sample size imbalance between preterm and term-born infants, we repeated the BAG analysis using a balanced subsample comprising 52 preterm infants and 52 matched term-born neonates (Fig. S22). Across all analyses, our findings remained highly consistent and robust, demonstrating minimal influence from variations in head motion thresholds, scan duration, sample size, and network resolution. These results confirm the reliability and stability of our conclusions across diverse analytical conditions.

## Discussion

Using personalized network modelling, we mapped developing cortical functional topography during the initial postnatal weeks. We found that while primary functional networks are already well established at birth, high-order association networks are present only as early prototypes. We observed that the refinement of these nascent networks occurs at their boundaries in an individualized manner and that this refinement process collectively predicts neonatal topography for both preterm and term-born infants, revealing altered maturational pattern in the association networks of preterm infants. These patterns of functional maturation are significantly correlated with indicators of concurrent structural development, particularly in terms of cortical myelination and sulcal depth. Together, our findings provide mechanistic insights into how the initial functional configuration of the neonatal brain provides a scaffold for the development of long-term cognitive ability.

The neonatal human brain already has complex neural configurations. At the cellular level, processes such as neurogenesis, synaptogenesis, and axon growth establish a foundational blueprint, priming the brain for postnatal axonal pruning and neural consolidation ^15, 54^. Functionally, persistent neuronal activity refines early neural circuits into organized spatial layouts (functional topography) ^40, 42, 43^ and reflects high postnatal plasticity ^55^. Building on this, we identified largely adult-like functional networks in the primary sensorimotor cortex, alongside still-maturing networks in the association cortex. Specifically, primary visual, somatomotor, and supplementary motor networks closely resembled their mature counterparts. In association networks, we delineated three fine-grained frontal networks ^30, 41^ and two distinct DMN components, moving beyond prior coarse descriptions ^30^. Notably, the emergence of high-order networks in neonates remains debated, with some studies reporting difficulty in detection ^41, 56^ and others confirming their presence at birth ^40, 53, 57^. Sylvester and colleague emphasized the importance of preserving long-distance connections to identify advanced functional networks ^53^, whereas others noted neonatal‒adult differences in cross-cortical connectivity ^58^. Our study demonstrates that effective machine learning algorithms such as NMF can recognize these networks at the individual level. Notably, these high-order networks, even in primordial forms, significantly predict 18-month neurodevelopmental outcomes at 18 months. For instance, the anterior DMN and lateral parietal network contribute to cognitive, language, and motor scores, underscoring their functional relevance from early development.

Interestingly, our results revealed a fragmented DMN organization in neonates, characterized by the separation of its anterior and posterior components. This contrasts with several neonatal studies reporting a more integrated adult-like network ^24, 53, 59^. A key methodological difference is that these prior studies employed surface-based mapping approach, whereas our framework was volume-based. Because surface-based mapping could globally generate more adult-like functional networks in neonatal scans, our fragmented DMN pattern should be interpreted with caution. In addition, differences in network decompose approaches and cohort heterogeneity may also contribute. For instance, even with surface-based mapping, separated neonatal DMN components emerge when connectivity threshold is altered ^53, 57^. Furthermore, using the same dHCP dataset, Williams and colleague identified an independent posterior-medial DMN component using a surface-based framework ^60^, providing support for our findings. Finally, we excluded subcortical regions because achieving highly reliable individualized mapping in these areas remains methodologically challenging. Subcortical nuclei are deep, small, and anatomically complex; thus, their BOLD signals are vulnerable to reduced receive sensitivity, partial-volume effects, physiological noise, and segmentation uncertainty ^61, 62^. Existing individualized subcortical mapping in adults relied on extensive resting/task-state pooling or dense within-subject sampling ^61–63^. These challenges are amplified in neonatal fMRI data by low deep-gray-matter SNR, small brain size, and short scan time ^64^. Nevertheless, the exclusion of subcortical regions may influence cortical component identification, particularly for cortico-subcortical systems such as the somatomotor network and DMN. Future studies employing resting/task-state integration ^63^, advanced 7T acquisitions ^65^, and multi-echo denoising ^66, 67^ will be crucial to address this issue.

We demonstrate that the fine-grained individual variation in neonatal function topography differs fundamentally between association networks and primary networks. While primary networks vary predominantly at their boundaries, association networks exhibit heterogeneity in both network cores and boundaries. This divergence likely stems from distinct cortical organizational principles and maturation dynamics ^30, 41^. Unlike the primary cortex, association networks are architecturally optimized for higher-order cognition through complex synaptic wiring, enhanced plasticity, and elevated metabolic demands ^68, 69^, potentially supported by fine-scale functional subnetworks with distributed representations ^70^. Developmentally, association networks remain in a dynamic state of spatial refinement during the neonatal period, shaped by genetic and environmental factors ^15, 71^, with substantial individual variation ^20–22^. In support of this, Xu *et al*. ^21^ reported that functional network variability decreases with age differentially between primary and association systems. Given the prolonged maturation of the association cortex into adolescence ^69^, future longitudinal studies integrating genetic and neuroimaging data across extended developmental windows will be crucial to elucidate these mechanisms.

The emerging “brain age” concept has proven valuable for characterizing typical and atypical brain maturation by quantifying the discrepancy between neuroimaging-predicted age and chronological age ^72, 73^. Previous studies have demonstrated that cortical functional connections, particularly in the sensorimotor cortex, can accurately predict an individual’s scan age ^74, 75^. Notably, we found that spatial refinement at network boundaries, especially in primary networks, contributes most substantially to brain age prediction, aligning with evidence that primary cortices undergo more pronounced early development than high-order regions do ^42, 76^. Histological studies have confirmed that primary cortices experience accelerated neurogenesis, synaptogenesis and axon growth around birth ^77^, supporting the emergence of core visual and motor functions ^54^. This early microstructural maturation may establish stable functional topography at network cores while maintaining developmental sensitivity at boundaries, creating a characteristic spatial maturation gradient. In our results, decreases in primary networks (Network 2 and 4) were accompanied by concurrent increases in association networks (Network 9 and 11). Age-related decreases have been widely reported across various functional metrics, including functional connectivity ^41, 76^ and local activity amplitude ^78^, in neonatal ^41, 76^, early-childhood ^79^, and adolescent ^9, 80^ cohorts. These reductions likely reflect increasing functional specialization, whereby functional representations become more spatially selective throughout development. Our findings capture functional recalibration along the cortical hierarchy axis ^69^ during neonatal stage. Prior studies have established prediction models in neonatal populations based on cortical morphology ^81–83^, white matter pathways ^84, 85^, and functional connectivity ^75, 85^. Our study extends these prior works by demonstrating that the individualized functional topography effectively captures early neurodevelopment ^81, 85^. Our approach achieves prediction accuracy comparable to functional connectivity-based models evaluated on the same dHCP dataset ^85^. Interestingly, models based on structural features generally yield higher prediction performance than those using functional measurements. This discrepancy likely arises because the underlying anatomical framework (e.g., cortical volume, folding, and myelination) is already largely established and undergoing fine-tuning during early in life ^15, 86^, whereas the cortical functional architecture remains nascent and highly dynamic ^18, 19^. Additionally, inherent confounders in neonatal functional imaging, such as sleep state, motion artifacts, and physiological noise, further limit the precision of individualized predictions ^73^. Finally, unlike most prior studies that trained BAG models separately on term or preterm samples, our model was trained exclusively on term-born neonates and directly applied to preterm infants at term-equivalent age. This design establishes a normative reference for typical development, enabling quantitative assessment of preterm deviations. However, direct cross-study comparisons on preterm BAG remain challenging due to the substantial methodological variations in model design.

Understanding the neurodevelopmental origins of diverse behavioural outcomes is critically important ^15, 87^. Our results demonstrate that individualized functional topography measured at birth accurately predicts cognitive, language, and motor abilities at 18 months of age. Several factors may explain this predictive relationship. First, despite its relatively coarse anatomical scaffold, the neonatal brain has already undergone substantial development ^54^, including neurogenesis, synaptogenesis, axonal growth, and initial myelination. These processes collectively establish the foundation for subsequent cognitive capabilities ^88, 89^. Second, although immature compared with adult benchmarks, neonatal functional configurations exhibit sufficient individual specificity to support identification ^90, 91^ and predict later learning ^90^ and cognition outcomes ^92^. Third, the early postnatal period represents a window of heightened plasticity during which brain function rapidly adapts to environmental inputs ^93^, enabling remarkable learning abilities even within hours of birth ^55^.

Notably, the most consistent predictive voxels for various neurodevelopmental outcomes clustered around network boundaries, particularly in primary sensorimotor networks, suggesting that these regions may function as common computational cores supporting diverse types of behavioural development. Previous research has indicated that compared with core regions, border zones of primary systems display distinct connectivity patterns, morphological features, and cytoarchitectonic organization ^94^. Functionally, these border areas predominantly occupy unimodal-to-limbic hierarchical positions ^94^, which are crucial for stabilizing and integrating inputs along the sensorimotor-to-association gradient ^94^. In our data, supplemental motor area boundaries showed prominent refinement—a region recently identified as part of an “action-mode network” implicated in action planning ^95^.

Our findings align with established developmental sequences: infants initially develop visual and motor abilities to support facial recognition, vocalization, and head control ^96^, which are abilities that foster secure attachment and facilitate higher-order skill acquisition ^97^. Moreover, association networks, including the DMN and lateral parietal cortex, differentially predicted specific 18-month outcomes, which is consistent with their known involvement in social cognition ^98^, rapid learning ^55^, and emotional processing ^99^. The mechanisms through which these networks support behavioural development are undoubtedly complex; future studies should integrate network communication models with task-based fMRI to elucidate how distributed neural interactions give rise to individual differences ^100^. Overall, the first two years of life represent a critical period for establishing lifelong behavioural foundations. Our findings provide potential neuroimaging biomarkers for tracking brain–behaviour relationships during this crucial window and may help elucidate the neural mechanisms underlying developmental neuropsychiatric disorders.

Preterm birth has been consistently linked to widespread neural alterations, including reduced cortical volume, microstructural changes, and disrupted white matter organization at term-equivalent age ^101, 102^. Functional abnormalities have also been reported in connectivity, network topology, and dynamics in preterm infants ^30, 103^. Extending these findings, we demonstrate that individualized functional topography reliably distinguishes preterm infants, with association networks contributing most significantly. These results align with dHCP-based reports of functional alterations in frontal‒parietal networks ^31^. Early exposure to extrauterine stimuli may place increased adaptive demands on developing association systems ^104^, which could bear the primary burden of accommodating this abrupt developmental transition. Interestingly, although prior studies describe marked abnormalities in primary visual or motor systems ^30, 47^, our approach revealed limited effects in these regions, possibly because the core spatial components of primary networks are already established and remain stable. We also reported altered functional maturation in preterm infants, which is consistent with reports of white matter ^105, 106^ and cortical development ^107^. Potential mechanisms include birth as an environmental trigger altering synaptic trajectories ^108^, early visual experience accelerating parvalbumin expression ^109^ and enhanced early nutrition. Finally, the model trained on term-born neonates generalized to preterm infants for brain age prediction, but not for neurodevelopmental outcomes. This dissociation suggests that core features of functional maturation are conserved across the two populations, whereas the link between neonatal functional organization and later behaviour is disrupted by preterm birth, potentially reflecting altered developmental trajectories and greater neurodevelopmental heterogeneity ^18^. Overall, our model enables the individualized quantification of this preterm maturation, offering a potential neuroimaging indicator of prematurity-related development.

Several potential limitations of this study should be acknowledged. First, the analysis was restricted to cortical regions. Given that subcortical areas and the cerebellum also undergo rapid structural and functional development during the neonatal period ^15^, future studies could incorporate these regions to examine how individualized functional topography evolves across the whole brain and its association with long-term neurocognitive outcomes. Second, previous research suggests that 18 months of age represents an early stage at which initial symptoms of certain neurodevelopmental disorders, such as autism, may develop ^110^. This underscores the importance of collecting longitudinal data from atypically developing populations to investigate how neonatal functional architecture influences the subsequent onset of neuropsychiatric conditions. Third, the relatively narrow age range of the current prediction model (spanning only 5 weeks of postnatal life) inherently limits its generalizability to broader developmental periods. Future work should further validate this framework in longitudinal neuroimaging and behavioural cohorts that encompass wider developmental windows, including early preterm and extended postnatal stages. Emerging large-scale developmental datasets, such as the BCP and HBCD ^111, 112^, alongside recent advances in ultra-high-field infant MRI ^65^, offer critical opportunities to establish a robust, generalizable BAG framework for infants and to further delineate its physiological and behavioural relevance. Finally, all neonatal fMRI scans in the dHCP dataset were acquired during natural sleep, a factor that must be considered when interpreting individual differences and developmental effects. Variations in sleep state may introduce biases into the estimation of functional connectivity and individual variability ^113–116^. Indeed, studies in older infants and toddlers have demonstrated that functional network organization differs between sleep and wakefulness ^114^. Furthermore, sleeping infants exhibit connectivity patterns that more closely resemble adult sleep than adult wakefulness ^115^. Although a recent study found largely preserved functional architecture across neonatal sleep stages ^116^, state-dependent effects cannot be entirely ruled out in our findings. Given that acquiring fMRI data from awake, healthy neonates remains highly impractical ^117^, future studies incorporating concurrent sleep-state monitoring will be crucial for disentangling true maturational effects from state-dependent variations in the neonatal brain.

In conclusion, we mapped personalized cortical functional topography during the first postnatal weeks, revealing that interindividual variability is already organized along a sensorimotor-to-association hierarchy. This spatial layout indicates the early emergence of adult-like variability patterns in the neonatal brain. Crucially, these individualized maps encode brain maturity in both term and preterm infants, predict early childhood neurodevelopmental outcomes, and are coupled with concurrent structural maturation. Our findings establish a foundational framework for the precision modeling of early brain development and its impact on long-term cognitive trajectories.

## Methods

### Participants and data acquisition

The whole dataset comprises a high spatial–temporal resolution of 367 term-born neonates and 52 preterm infants as part of the Developing Human Connectome Project ^35^. All infants were scanned during natural sleep without sedation to obtain structural and resting-state fMRI data using a 3T Philips Achieva scanner. Bayley-III scores were assessed in 249 infants at 18 months of age to capture individual developmental outcomes in motor, language, and cognitive domains ^46^. The Q-CHAT score for 245 infants at 18 months of age was also assessed to measure atypical social, sensory, and repetitive behaviour risks ^118^. Written consent was obtained from the guardian of each baby. The dataset was further stratified into three groups: (1) Subset 1 included 52 term-born neonates (36 females; GA at birth, 37–42 weeks; PMA at scan, 37–44 weeks, 15 minutes scan). This dataset was used to construct a typical neonatal group atlas. (2) Subset 2 included the remaining 315 term-born neonates (134 females; GA at birth, 37–42 weeks; PMA at scan, 37–44 weeks, 15 minutes scan) and was used to reconstruct individualized functional topography to delineate functional variability and to conduct prediction analysis on brain maturity and neurocognitive outcomes. (3) Subset 3 included 52 preterm infants (19 females; GA at birth, 23–36 weeks; PMA at scan, 37–44 weeks, 15 minutes scan) and was used to conduct classification analysis between preterm and term infants. Notably, Subset 1 was derived from dHCP Release 2, whereas Subsets 2 and 3 were selected from dHCP Release 3. Identical quality control and preprocessing procedures were applied across all datasets. Importantly, data from Subset 1 were strictly excluded from the personalized topographic analyses. This data-splitting strategy prevents circularity and ensures unbiased estimation for all subsequent individual-level analyses. Details of datasets are provided in Table 1.

For comparison purposes, we also included 200 adults from the public HCP S1200 dataset ^37^ (73 females, 22–25 years old, 14 minutes scan). We selected the brain scans of the 200 youngest healthy adults (73 females; ages 22–25 years; 14-minute scans) from all the participants. Written informed consent was obtained from all the subjects, and the scanning protocol was approved by the Institutional Review Board of Washington University in St. Louis, MO, USA (IRB #20120436). Details are provided in Supplementary Section 1.

### MRI data preprocessing

For all the fMRI images in the dHCP dataset, we performed three preprocessing stages: minimal preprocessing, customized spatial normalization, and final-stage preprocessing. The minimal preprocessing steps were performed by the dHCP organizer and have been validated in prior studies ^119^. To obtain good spatial correspondence, spatial normalization steps were designed by generating a customized template with a Serag template ^120^ at 37 weeks initially using all the brain scans from Subset 1. Individual T2w images and tissue masks were nonlinearly registered to the customized template. Final-stage preprocessing was performed using GRETNA v2.0.0, and included smoothing, linear detrending, nuisance variable regression, scrubbing, and temporal filtering. Notably, Friston’s 24 head motion parameters and average signals were removed using multivariable linear regression, and spike regression-based scrubbing (0.5 mm displacement) ^121, 122^ was performed to control for the effect of head motion. For frames control, individual scans were retained only if they contained at least 70% clean volumes of all time points. For the HCP dataset, all functional images initially underwent minimal preprocessing using the HCP public pipeline ^123^. Following this, similar processing steps as those for the dHCP dataset were performed.

#### Generating individualized brain networks

##### 1) Group network identification

To make a typical representation of the broadly consistent functional network across the neonatal stage, we first generated a group network atlas, which was then used as initialization for subsequent individualized network generation. As previously described ^9, 36^, we used NMF to derive these group networks. The NMF approach employs a group consensus regularization term that preserves the interindividual correspondence of labels and locality between functional networks and ensures that the decomposition is robust to imaging noise (see ^36^ for details). We set the number of networks as 11 ^19, 30^ in the main analyses and validated it at 9 and 17. To reduce the influence of outliers, a bootstrap strategy was used to perform group-level decomposition on 80% of the subjects randomly selected each time, which was repeated 50 times. During each iteration, a time series matrix with 92 000 rows (time points) and 6905 columns (voxels) was constructed, and a network loading matrix was constructed, which yielded 50 different group atlases in total. We combined these atlases into a robust loading matrix V using spectral clustering ^36^. The resulting group-level network loading matrix V had N rows (network number) and 6905 columns (voxel number), representing the loading of each given voxel.

##### 2) Individualized network mapping

We mapped each individualized specific network atlas using a second-time regularized NMF based on the identified group networks as initialization and individual fMRI time series (2300 × 6905) as input, which yielded a loading matrix V (11 × 6905 matrix) for each participant. This probabilistic (soft) definition was converted into discrete (hard) network definitions for display by labelling each voxel according to its highest loading.

#### Individual variation in neonatal functional network topography

To represent the individual variation in functional network topography, we first computed the voxel-wise variability by calculating the median absolute deviation of the probability network loading across neonates for each voxel and summing across networks to generate a whole-brain variability map. Group differences between primary and association networks were assessed using spin-based permutation testing.

Next, four network-level topographic properties were evaluated: 1) position variability—we calculated the average Euclidean distance among the parcel centres (centre of mass) across subjects; 2) size variability—we calculated the standard deviation of size (the number of voxels of a region) across subjects; 3) overlapping variability—we calculated the mean Dice coefficient deviation between each pair of subjects; and 4) homogeneity variability—we calculated the standard deviation of homogeneity across subjects (Fig. S4; for a detailed algorithmic workflow, please refer to ^45^). All comparisons between primary and association networks were tested using 10,000 spin permutations.

#### Null model testing

Two null models were employed for statistical analyses. First, to evaluate the significance of the prediction models, we performed permutation testing (*P*_perm_). Prediction labels were randomly shuffled 1,000 times, and the models were retrained to generate a null distribution of prediction accuracies. 2) To determine whether cortical system differences were driven by spatial autocorrelation ^124^, we applied a spin test (*P*_spin_). Cortical maps were randomly rotated 10,000 times to generate spatially autocorrelated surrogate Z-maps, providing a null distribution for the observed system-level differences.

#### Associations of network topography with development, neurodevelopmental outcomes at 18 months, and preterm birth

Given the nonlinear nature of brain development, we used GAMs to capture the associations between network topography and scan age while controlling for sex, mean FD, and scan-birth age intervals. To evaluate 18-month neurodevelopmental outcome associations, GAMs were similarly fitted with outcome scores as dependent variables, adjusting for sex, mean FD, scan age, and the time interval between birth and the scan. To evaluate preterm birth associations, preterm status (preterm vs. term) was included as the primary group factor, with sex, mean FD, and scan age as covariates. These models were fitted at both the network and voxel levels. At the network level, FDR correction was employed to control for family-wise error. At the voxel level, the significance threshold was set at *P* < 0.001 at the voxel level, with Gaussian random field (GRF) correction at the cluster level of *P* < 0.05 ^125^.

#### Multivariable topographic network-level prediction framework (TiPCA model)

To assess whether and how individualized functional network topography predicts brain maturity and neurodevelopmental outcomes and to classify preterm infants from their term-born counterparts, we developed a multivariable TiPCA prediction model approach based on the SVR framework. This approach involves performing feature reduction by separately performing PCA within each network before model training. Within this framework, feature dimensionality was reduced by applying PCA separately within each functional network prior to model training, retaining (N − 1) components per network, where N denotes the number of training samples. For the SVR model, the linear kernel was employed. We validated the robustness of the prediction model by testing prediction accuracy using repeated random 10F-CV, repeating 10F-CV 100 times and averaging the correlation to determine overall prediction accuracy. To identify high-contributing networks, we evaluated both contributing weights and cross-fold stability. For each network, we defined a contribution score as the product of its mean weight and mean rank across all random folds. The resulting scores were then z-transformed, and networks with positive z-scores were classified as high-contributing. To further depict the spatial topography of high-contributing voxels, we calculated the voxel-wise feature contribution weights within the high-contributing network and summed the absolute weights across networks.

The TiPCA framework was consistently used for the prediction of individual scan age, Bayley-III scores and Q-CHAT scores with a 10-fold cross-validation strategy. Different confounding variables were considered for each prediction. To predict brain maturity, we regressed out the confounding variables of sex, mean FD, and the time interval between birth and the scan from both functional topography and scan ages. To predict behavioural assessments, we regressed out confounding variables (age at scan, sex, mean FD, and scan-birth age intervals) from both functional topography and neurodevelopmental outcomes.

#### Cortical structural maturation and structure‒function association analysis

To investigate whether neonatal functional topographic refinement is constrained by cortical structural maturation, we employed structural MRI scans from 301 term-born neonates (Subset 2). For each participant, we derived individual cortical surface maps for four anatomical features: cortical myelination (T1w/T2w ratio), sulcal depth, cortical thickness and surface curvature. We assessed age-related changes in each structural feature using vertex-wise GAMs, with scan age as the main factor and sex, mean FD, and the interval between birth and scan age as covariates. The resulting F-statistic maps quantified the spatial pattern of structural maturation across the cortex. Significance was determined using vertex-wise FDR correction at *q* < 0.05.

To examine structure‒function relationships, we performed PLS analyses between the structural maturation maps (*F* values) and the contribution weights of functional topography derived from predictive models of brain age and behavioural outcomes. Specifically, we computed spatial correlations between functional feature weights (from the TiPCA model) and the structural maturation maps in the PLS model at both the global and network levels. We summarized the PLS loadings across the four structural features to identify the dominant contributors. FDR correction was applied across networks.

#### Identifying preterm infants and estimating personalized brain age gaps

To classify term-born and preterm infants at term-equivalent age, we employed a similar TiPCA classification approach with a 10-fold cross-validation strategy using Subset 2 and Subset 3 (the 52 preterm infants and matched 73 term neonates (scan age, sex, and head motion matched)). Before classification, we regressed out confounding variables (age at scan, sex, and mean FD) from functional topography. We then evaluated the network-wise feature weights for preterm and term classification to determine their contributions to the model.

To quantitatively assess the degree of abnormal maturation in preterm infants, we used preexisting brain maturity prediction models trained on term-born neonates to predict the PMA at scan for each preterm-born infant as the brain age. By calculating the difference between the predicted postmenstrual age and the actual postmenstrual age, we derived the personalized brain age gap (BAG) for each preterm infant. To adjust for prediction biases ^48^, we corrected the slope and intercept of BAG (Fig. S14). Specifically, within the training set, predicted age was regressed on chronological age to estimate the intercept and slope of the bias. The resulting parameters were used to adjust the predicted ages of the test samples. The BAG was subsequently calculated as the adjusted predicted age minus chronological age. To determine whether brain maturation is altered in preterm infants relative to term-born infants, we applied a two-sample t test to compare the BAG between the two groups. We also examined associations between the BAG and 18-month neurodevelopmental outcomes using partial correlations while controlling for sex, scan age, FD, and the time interval between birth and the scan.

Finally, we explored whether alterations in functional topography in preterm infants were constrained by structural abnormalities. Individual cortical maps of myelination, thickness, curvature, and sulcal depth were derived and aligned to a common surface template. Group differences were assessed using GAMs, with sex, mean FD, and scan age as covariates. Statistical significance was determined using FDR correction (*q* < 0.05). A similar PLS analysis was used to test the association between group-level structural abnormalities (Z statistics of group differences) and network-wise contribution weights from the classification model.

#### Sensitivity analysis

We conducted multiple sensitivity analyses to ensure the robustness of our findings. Specifically, we (i) repeated all analyses using a stricter head motion threshold (mean FD < 0.3 mm); (ii) evaluated the effect of scan-duration on the stability of functional topographies using a split-half analysis with both sequential and random sampling frames (1–7.5 min) ^24, 126^; (iii) evaluated the effect of sample size on brain age and brain–behaviour predictions through bootstrapped subsampling across logarithmically spaced sample sizes; (iv) performed split-half validation by dividing neonates into two age-matched subsets; (v) regenerated group-level atlases using randomly selected subsets of term-born neonates (n = 10–300); (vi) evaluated the impact of network resolutions using alternative parcellations comprising 9 and 17 networks; (vii) assessed the effect of estimating covariates within training folds versus across the full sample; and (viii) investigated potential biases arising from imbalanced sample sizes between preterm and term-born infants. Further methodological details are provided in Supplementary Information, Section 6.

## Data availability

All data required for reproducing our findings have been publicly available, including the group-level network, the individualized functional networks, individual variability, brain maturity prediction model, neurodevelopmental outcomes prediction model, structure–function coupling, preterm functional topographic alterations, and the data for visualizing main figures. They are stored in a publicly accessible cloud repository (https://github.com/zhaohuaxishi1/neonatal_Individual_Parcellation). For the dHCP dataset, raw image scans are available under restricted access in accordance with the dHCP data release conditions via https://data.developingconnectome.org. For the HCP dataset, raw image scans are publicly available through ConnectomeDB: https://db.humanconnectome.org/app/template/Login.vm. Source data are provided with this paper.

## Code availability

The codes used in this study ^127^ are available at GitHub: https://github.com/zhaohuaxishi1/neonatal_Individual_Parcellation. Software packages used in this manuscript include dHCP structural pipeline v1 (https://github.com/BioMedIA/dhcp-structural-pipeline), dHCP functional pipeline v1 (https://git.fmrib.ox.ac.uk/seanf/dhcp-neonatal-fmri-pipeline), SPM 12 (http://www.fil.ion.ucl.ac.uk/spm/), GRETNA toolbox 2.0.0 (https://www.nitrc.org/projects/gretna), Individual Parcellation method (https://github.com/hmlicas/Collaborative_Brain_Decomposition), BrainSMASH (https://github.com/murraylab/brainsmash/), LIBSVM (3.25) (https://www.csie.ntu.edu.tw/~cjlin/libsvm/), Support Vector Regression (https://github.com/ZaixuCui/Pattern_Regression_Clean), Support Vector Classification(https://github.com/ZaixuCui/Pattern_Classification), cifti-matlab toolbox v2 (https://github.com/Washington-University/cifti-matlab), BrainNet Viewer 1.7 (https://www.nitrc.org/projects/bnv), R 4.0.3 (https://www.r-project.org), Matlab 2020b (https://www.mathworks.com/products/matlab.html), and Python 3.7.0 (https://www.python.org), which are available online.

## Acknowledgments

This work was supported by the National Natural Science Foundation of China (Nos. 82672670 (T.Z.), 82021004 (Y.H.), 31830034 (Y.H.), 82102131 (Y.X.)), the Beijing Natural Science Foundation (No. 7262077 (T.Z.)) and the Fundamental Research Funds for the Central Universities (Nos. 2233300002 (T.Z.), 2233100018 (T.Z.)).

## Author Contributions

J.L.Z., T.D.Z. and Y.H. designed research; T.D.Z., Y.H.X., H.M.L., Q.L.L., L.L.S., X.Y.L., M.Z.H., Z.L.Z. and Z.X.C. provided the methodological instruction; J.L.Z. and T.D.Z. performed the data analysis; J.L.Z. and T.D.Z. developed visualizations. J.L.Z., T.D.Z., and Y.H. wrote the paper; J.L.Z., T.D.Z., and Y.H. revised the paper.

## Conflicting Interests

The authors have declared that no conflicting interests exist.

## Notes

https://github.com/zhaohuaxishi1/neonatal_Individual_Parcellation

### Competing Interest Statement

The authors have declared no competing interest.

### Summary of Updates

This version has been substantially revised following peer review. We updated the manuscript text and figures, clarified the methods and statistical reporting, added additional validation and sensitivity analyses, and revised the supplementary materials. The main conclusions of the study remain unchanged.

