## Supplementary Information for "Precision probabilistic mapping reveals personalized functional topography in the early postnatal human brain"

### SI-1. Participants and MRI Data Analyses

#### 1.1 Participants

The neonatal data were obtained from the Developing Human Connectome Project (dHCP) [1] (http://www.developingconnectome.org/), with ethical approval (14/LO/1169) and informed parental consent. Neurodevelopmental outcomes at 18 months were assessed using the Bayley Scales of Infant and Toddler Development, Third Edition (Bayley-III) [2]. The Quantitative Checklist for Autism in Toddlers (Q-CHAT) score at 18 months age was also assessed to measure the atypical social, sensory, and repetitive behavior risks [3]. Following stringent quality control procedures (see details in Fig S1), 367 term-born (170 females; gestational age [GA] at birth = 37-42 weeks; postmenstrual age [PMA] at scan = 37-44 weeks; scan duration = 15 minutes) and 52 preterm infants were included. Participants were stratified into three subsets for specific analyses: (i) Subset1: 52 term-born neonates from dHCP Release 2 (36 females; GA at birth, 37–42 weeks; PMA at scan, 37–44 weeks, 15 minutes' scan), which was utilized to construct a typical group atlas. (ii) Subset 2: 315 term-born neonates from dHCP release 3 (134 females; GA at birth, 37–42 weeks; PMA at scan, 37–44 weeks, 15 minutes' scan), used for individual functional networks reconstruction, variability analysis, prediction of brain maturity and 18-month neurodevelopment, and structural basis of the early functional topographic development. (iii) Subset 3: 52 preterm infants (19 females; GA at birth, 23–36 weeks; PMA at scan, 37–44 weeks, 15 minutes' scan), used to assess alteration of functional topography in preterm infants. To compare neonatal and adult functional variability patterns, we additionally analyzed data from the Human Connectome Project (HCP) [4]. For detailed image inclusion/exclusion criteria, please refer to [4]. From this dataset, we selected 200 healthy young adults (73 females; age = 22-25 years; scan duration = 14 minutes) with minimally preprocessed resting-state fMRI data. Informed consent was obtained from all participants, and all procedures were approved by the Institutional Review Board at Washington University in St. Louis (IRB #201204036).

#### 1.2 Image Acquisition

For the dHCP dataset, resting-state fMRI and structural MR scans were acquired for each neonate during natural sleep without sedation, using a 3T Philips Achieva scanner with a 32-channel neonatal coil. Full details regarding the data collection have been described [5]. BOLD functional MRI with high spatial and temporal resolution were scanned using the following parameters: repetition time (TR) = 392 ms, echo time (TE) = 38 ms, flip angle (FA) = 34°, voxel size = 2.15 × 2.15 × 2.15 $mm^3$, number of volumes = 2300, multiband factor = 9, scan time = 15 min 3 s. To correct the susceptibility artifacts for BOLD images, inversed encoding spin-echo field maps were obtained. High-resolution T1- and T2-weighted MR images were acquired with following parameters: T1-weighted: TR = 4795 ms, TE= 8.7 ms, 0.8 mm isotropic voxels, field of view (FOV) = 145 × 122 × 100 mm, T2-weighted: TR = 12s, TE = 156 ms, 0.8 mm isotropic voxels, FOV = 145 × 145 × 108 mm.

For the HCP dataset, all MRI data were acquired on a customized 3T 32-channel Siemens Skyra scanner at Washington University. resting-state fMRI images were obtained by gradient-echo-planar imaging acquisitions with two separate runs. The sequence parameters for each run were the same as follows: TR = 720 ms, TE = 33.1 ms, FA = 52°, bandwidth = 2290 Hz/pixel, FOV = 208 × 180 mm, 72 slices, voxel size = 2 × 2 × 2 $mm^3$, multiband factor = 8, and scan time = 14 min 33 s. T1- and T2-weighted MR images were acquired with following parameters. T1-weighted: TR = 2400 ms, TE = 2.14 ms, 0.7 mm isotropic voxels, matrix = 320 × 320, 256 slices, FA = 8°, FOV = 224 × 224 mm.T2-weighted: TR = 3200 ms, TE = 565 ms, 0.7 mm isotropic voxels, FOV = 224 × 224 mm.

### SI-2. Generating individual functional networks and quantifying interindividual variability

#### 2.1 Group-level and individualized network generation

As the same approach in previous studies on adult and adolescent brain [6, 7], we employed the nonnegative matrix factorization (NMF) framework to derive individualized functional networks[8]. This approach employed a group consensus regularization term that preserves the inter-individual correspondence, as well as a data locality regularization term that makes the decomposition robust to imaging noise (see [6] for details of the method; see also: https://github.com/hmlicas/Collaborative_Brain_Decomposition). As NMF requires the input to be nonnegative values, we rescaled the data by shifting the time courses of each voxel linearly to ensure all values were positive[6] and normalized the time course by its maximum value to ensure all input values in the range of [0, 1]. To obtain individualized networks, two stages of NMF iterations were employed as follows.

Given a group of n participants, each having fMRI data $X^i \in R, i = 1, \dots, n$, consisting of $S$ voxels and $T$ time points, we aimed to find K non-negative functional networks $V^i = \left(V^i_{s,k}\right) \in R^{S\times k}$ and their corresponding time courses $U^i = \left(U^i_{t,k}\right) \in R^{T\times K}$ for each neonate, such that

$$X^i \approx U^i\left(V^i\right)' + E^i, s.t. U^i, V^i \geq 0, \forall 1 \leq i \leq n,$$

where $(V^i)'$ is the transpose of $(V^i)$, and $E^i$ is independently and identically distributed (i.i.d) residual noise following Gaussian distribution with a probability density function of $g(x) = \left(\frac{1}{\sqrt{2\pi}\sigma}\right) e^{-\frac{x^2}{2\sigma^2}}$. Both $U^i$ and $V^i$ were constrained to be non-negative to guarantee each functional network contains no anti-correlated functional units [8]. A group consensus regularization term was applied to ensure inter-individual correspondence, which was implemented as a scale-invariant group sparsity term on each column of $V^i$ and formulated as

$$R_c = \sum_{k=1}^{K} \left\| V^{\widetilde{1,\dots,n}}_{.,k} \right\|_{2,1} = \sum_{k=1}^{k} \frac{\sum_{s=1}^{S}\left(\sum_{i=1}^{n}\left(V^i_{(s,k)}\right)^2\right)^{\frac{1}{2}}}{\left(\sum_{s=1}^{S}\sum_{i=1}^{n}\left(V^i_{s,k}\right)^2\right)^{\frac{1}{2}}}$$

The data locality regularization term was applied to encourage spatial smoothness and coherence of the functional networks using graph regularization techniques[9]. The data locality regularization term was formulated as

$$R^i_M = T_r\left(\left(V^i\right)' L^i_M V^i\right),$$

where $L^i_M = D^i_M - W^i_M$ is a Laplacian matrix for participant I, $W^i_M$ is a pairwise affinity matrix to measure spatial closeness or functional similarity between different voxels, and $D^i_M$ is its corresponding degree matrix. The similarity between each pair of spatially connected voxels (i.e., voxels a and b) was calculated as $(1 + corr(X^i_{.,a}, X^i_{.,b}))/2$, where $corr(X^i_{.,a}, X^i_{.,b})$ is the Pearson correlation coefficient between time series $X^i_{.,a}$ and $X^i_{.,b}$, and others were set to zero so

that $W_M^i$ has a sparse structure.
We identified individual functional networks by optimizing a joint model with integrated data fitting and regularization terms formulated by

$$\min_{(U^i,V^i)} \sum_{i=1}^{n} \left\| X^i - U^i(V^i) \right\|_F^2 + \lambda_M \sum_{i=1}^{n} R_M^i + \lambda_C R_C,$$

$$s.t. U^i, V^i \geq 0, \left\| V_{\cdot,k}^i \right\|_\infty = 1, \forall 1 \leq k \leq K, \forall 1 \leq i \leq n$$

where $\lambda_M = \beta \times \frac{T}{K \times n_M}$ and $\lambda_C = \alpha \cdot \frac{n \cdot T}{K}$ are used to balance the data fitting, data locality, and group consensus regularization terms, $n_M$ is the number of neighboring voxels, $\alpha$ and $\beta$ are free parameters. Of note, we used identical parameters settings as in prior studies on individualized functional network mapping [6].

#### 2.2 Comparison of functional homogeneity between individualized networks and group-level networks

Network homogeneity is a commonly used metric for evaluating the success of functional parcellation [10-12]. As previously described, network homogeneity was calculated as the average of Pearson's correlation coefficients between the time series of all pairs of voxels within each network [11]. To summarize network homogeneity for comparisons of group networks and individual networks, we averaged the homogeneity value across networks.

#### 2.3 Estimating the individual variability in topographic properties at the network level

We calculated four indices to separately describe the position, size, overlap, and regional homogeneity of each network and assessed the individual variability of each indicator. The definitions of these indices are as follows.

The position of a functional region was represented by the coordinates of its centre of mass. For each parcel, the interparticipant variability in parcel position was estimated as the average Euclidean distance among the parcel centres across participants, normalized by the maximum average Euclidean distance. Specifically, the individual variability in the position of a functional network *i* was defined as follows:

$$V_i = \frac{E[\sqrt{\left(F_i(S_p) - F_i(S_q)\right)^2}]}{max[\sqrt{\left(F_i(S_p) - F_i(S_q)\right)^2}]}$$

where $p$, $q$ = 1,2…$M$ (p ≠ q), $M$ is the number of infants, and $F_i(S_p)$ is the position of a functional network *i* in infant *p*.

The size of a functional region was calculated as the number of voxels that fell within that region. For each parcel, the interparticipant variability in size was calculated as the standard deviation of size across participants. Specifically, the individual variability in the size of a functional network i was defined as follows:

$$V_i = \mathrm{E}[\frac{|F_i(S_p) - F_i(S_q)|}{\max\left(F_i(S_p), F_i(S_q)\right)}]$$

where $p$, $q$ = 1,2…$M$ ($p \neq q$), $M$ is the number of infants, and $F_i(S_p)$ is the size of a functional network $i$ in infant $p$.

The overlap of a functional network was calculated as the Dice coefficient between the individualized area and group-level parcel. For each network, the interparticipant variability of overlap was calculated as 1 minus the mean Dice coefficient deviation across all pairs of participants. Specifically, the individual variability in the overlap of a functional network i was defined as follows:

$$V_i = 1 - \mathrm{E}[\frac{2|F_i(S_p) \cap F_i(S_q)|}{|F_i(S_p)|+|F_i(S_q)|}]$$

where $p$, $q = 1,2\ldots M$ ($p \neq q$), $M$ is the number of infants, and $F_i(S_p)$ is the functional network i in infant p.

The homogeneity of a functional network was calculated as the mean homogeneity within all voxels. For each parcel, interparticipant variability in parcel homogeneity was calculated as the standard deviation of homogeneity across participants, normalized by the maximum deviation. Specifically, the individual variability of the homogeneity of a functional network i was defined as follows:

$$V_i = \mathrm{E}[\frac{|F_i(S_p) - F_i(S_q)|}{max\left(F_i(S_p), F_i(S_q)\right)}]$$

where $p$, $q = 1,2\ldots M$ ($p \neq q$), $M$ is the number of infants, and $F_i(S_p)$ is the mean homogeneity of a functional network i in infant p.

#### 2.4 Comparison of functional topographic variability between neonates and adults

Like in previous adult studies [13, 14], we set the number of functional networks to 17 and identified all functional networks by the same NMF procedure. We computed voxel-wise individual variability maps using the same definition as that used for the neonatal brain. To compare the patterns of the functional variability maps between neonatal and adult brains, the adult variability map was linearly registered to the neonatal brain using the antsRegistrationSyN method [15].

### SI-3. Prediction of brain maturity and neurodevelopmental outcomes at 18 months from functional network topography

#### 3.1 Multivariable prediction using topographic network-level modelling (Ti-PCA model)

To examine whether individualized functional network topography predicts brain maturity and neurodevelopmental outcomes, we established a Ti-PCA model using linear support vector regression (SVR). Specifically, we performed network-wise principal component analysis (PCA) on the voxel-wise functional topographies separately for each network. The low-dimensional features from all the networks were then concatenated into a linear SVR prediction model (Fig. S6). This approach preserved the independence of topographic features from each network during feature dimension reduction.

The models were trained and tested with a 10-fold cross-validation approach (10F-CV). To predict brain maturation, 315 term-born neonates (134 females; 41.35 ± 1.68 PMA at scan) in Subset 2 were used. Confounding variables, including sex, mean framewise displacement (FD), and the time interval between birth and the scan, were regressed out from both the input features and the prediction targets prior to modelling. For behavioural predictions, 249 neonates (135 males; 41.39 ± 1.7 PMA at scan and 19.1 ± 2.2 months at follow-up) had valid neurodevelopmental assessments, including cognitive, language, and motor scores, at 18 months. Age at the scan was additionally controlled for as a confounding variable. The regularization parameter C was set to 1, following previous recommendations [16].

##### 3.1.1 10F-CV and random splitting 100 times

To minimize the overfitting effect while ensuring enough training samples, we employed a 10F-CV approach by dividing all the participants into 10 subsets. Specifically, we sorted the participants according to the predictors (i.e., age or neurodevelopmental outcomes at 18 months) [17]. We iteratively used each subset as the testing set, with the other subsets used as the training set. Each feature was linearly normalized such that it had zero mean and unit variance across the training dataset, and the normalization parameters were also applied to normalize the testing dataset. Notably, network-wise PCA was performed on the training set, retaining N-1 components per network (N = number of training participants), and the same reduction coefficients were applied to the testing set. Prediction accuracy was quantified as the Pearson correlation between predicted and actual values across all folds. Here, we used the LIBSVM function in MATLAB to implement support vector regression (https://www.csie.ntu.edu.tw/~cjlin/libsvm/;[18]).

To avoid the arbitrary splitting of 10 subsets, we also repeated the grouping of subsets randomly 100 times and obtained a distribution of prediction accuracy to represent the model performance.

##### 3.1.2 Significance of prediction performance

To evaluate whether the prediction performance (i.e., correlation r) was significantly better than expected by chance, we performed a permutation test [19]. Specifically, we permuted the prediction labels (i.e., age or neurodevelopmental outcomes at 18 months) across the training samples randomly 1,000 times (both retrained and retested). The p value was calculated for the proportion of permutations that exhibited a higher value than the actual r value of the real label.

##### 3.1.3 Interpretation of model feature weights at both the network and voxel levels

Our Ti-PCA framework enables network-specific contributions to be estimated without feature mixing during model training. However, because all network-level features are simultaneously entered into the model, the multicollinearity effect may obscure the contribution of certain networks. To address this, we separately entered the functional topography of each network into the SVR model and further identified high-contributing networks by jointly considering contributing weights and ross-fold stability across 100 random splits. For each network, we defined a contribution score as the product of its mean weight and mean rank across all random splits. The resulting scores were then z-transformed, and networks with positive z-scores were classified as high-contributing.

To characterize the voxel-wise distribution of high contributing features, we averaged the voxel-wise contribution weights across all the networks and retained only the contributing voxels of the high-contributing networks, yielding the final contribution map. To assess how voxel-wise feature contributions are related to network loading and interindividual variability, voxel-wise contribution weights were correlated with network loading and interindividual variability. To further quantify the relative influence of each network on these correlations, we performed a correlation decomposition analysis, in which the mean contribution value within each network was considered an index of its contribution magnitude.

### SI-4. Structural basis of functional topographies contributing to typical and atypical brain maturation and behavioural outcomes

#### 4.1 Structural cortical features

Structural cortical features were obtained from dHCP Release 3 (http://www.developingconnectome.org/) for 301 term-born and 41 preterm infants separately in Subset 2 and Subset 3 provided by the dHCP structural pipeline[20]. We employed four anatomical features that are well-established neuroimaging markers of structural changes during early postnatal brain development, including cortical myelination (T1w/T2w ratio), thickness, curvature, and sulcal depth.

#### 4.2 Mapping of structural features onto the fsaverage4 surface template

The individual vertex-level structural metrics were first projected to the 40-week PMA template using multimodal surface matching (MSM), optimized for the alignment of sulcal depth features provided by the dHCP pipeline [21, 22]. The 40-week template was subsequently registered to the HCP fs_LR_32k template using MSM provided by Logan Z. J. Williams [21]. Final resampling to the fsaverage4 standard mesh (2,562 vertices per hemisphere) was performed using the HCP Workbench metric-resample command [23], enabling cross-participant comparisons in a common cortical surface space.

#### 4.3 Mapping of functional features onto the fsaverage4 surface template

We mapped our neonatal functional features onto the fsaverage4 surface template. The registration included two stages: first, nonlinear transformations from the neonatal T2 template space to the MNI space using the antsRegistrationSyN method [15] and then nonlinear projections from the MNI volume space to the fsaverage 4 space using the bbregister method [24].

#### 4.4 Structural changes underlying neonatal functional topographic refinement

We first quantified the maturation of each anatomical feature using vertex-wise GAMs, with scan age as the primary effect and sex, mean FD, and the interval between birth and scan as covariates. The F-statistic of the brain maps was calculated to represent the spatial distribution of the degree of maturation (vertex-wise FDR correction at $q < 0.05$).

#### 4.5 Partial least squares (PLS) to identify structural constraints on the contribution weights of functional topography

We performed PLS analyses between the structural maturation maps (F values) and the functional contribution maps derived from the predictive models (brain age and behavioural outcomes). Such analyses were performed at both the whole-brain and network levels. The PLS loadings were summarized within each of the four structural features to identify dominant contributors. Bonferroni correction was applied across networks.

The same analysis was also performed for preterm infants. The group differences in the four anatomical features between 41 preterm and 56 term-born infants were assessed using vertex-wise GAMs, with sex, scan age, and mean FD as covariates. The Z-statistic map was calculated for each feature, and statistical significance was determined using vertex-wise FDR correction ($q < 0.05$). We then examined whether the maps representing the structural abnormalities of preterm infants were associated with the functional contribution map from the Ti-PCA classification model (term vs. preterm). PLS analyses were performed to estimate structure–function associations. Such analyses were performed at both the whole-brain and network levels. The PLS loadings were taken as their absolute values and then summarized to identify the dominant contributors among the four structural features. Bonferroni correction was applied across the networks.

### SI-5. Classification and brain maturation deviation analysis of the preterm brain from functional network topography

#### 5.1 Using Ti-PCA classification modelling to distinguish preterm infants from term-born neonates on the basis of functional network topography

In Subset 3, we employed the data of 52 preterm infants and 73 matched term-born neonates (scan age, sex, and head motion matched) to assess whether the multivariable spatial pattern of network topography could distinguish preterm infants from term-born neonates. We adopted a multi-step matching approach to maximize distributional similarity between groups: 1) **Age matching.** all term-born infants were first stratified into four PMA bins (37–39, 39–41, 41–43, and 43–44 weeks) to match the scan-age distribution of the preterm cohort. 2) **Head motion matching.** Within each PMA bin, we selected term-born infants whose mean FD (head-motion degree) fell within ± 0.5 standard deviations of the mean value in the corresponding preterm group. 3) **Sex matching.** Within each PMA bin, we selected female term-born infants satisfying the same FD criteria until the sex proportion was matched to that of the preterm group. After these, 73 term-born infants were finally selected to provide the closest overall match to the preterm cohort.

To address this question, we used a similar Ti-PCA classification approach with 10F-CV to unbiasedly classify preterm and term-born neonates. Before classification, the confounding variables of age at scan time, sex, and mean FD were regressed out from the functional topography. The prediction model was evaluated using 10-fold cross-validation with random splits. The statistical significance of the classification accuracy was assessed using permutation tests with 1000 random resamples of labels.

#### 5.2 Estimating individualized deviation (brain age gap) of the preterm brain from normative age prediction models

Recent progress in neuroimaging studies has shown that the brain age gap (BAG) may serve as a promising biomarker for capturing individual deviations in brain maturation and indicating potential risks of neurodevelopmental disorders [25, 26].

For each infant, the BAG was calculated by subtracting the actual PMA at the scan from the predicted PMA. To account for regression-to-the-mean bias [27], we corrected the slope and intercept of the predicted age vs. actual age fitting (Fig. S14). On the basis of the preestablished brain maturity prediction model derived from individualized functional topography in 315 term babies, BAGs were computed at both the whole-brain (loading maps of all 11 networks) and network (the loading map of one of the 11 networks) levels for 315 term-born and 52 preterm infants.

To determine whether the brain maturation of preterm infants is delayed or accelerated relative to that of term-born controls, we compared the BAG between term-born neonates and preterm infants using two-sample t tests at both the whole-brain and network levels. During the comparison, we focused only on metrics that significantly predict brain age for both groups.

Finally, we assessed correlations between the BAG and 18-month neurodevelopmental outcomes (cognitive, language, and motor scores), controlling for sex, scan age, mean FD, and the scan–birth interval.

### SI-6. Sensitivity analysis

We performed systematic validation analyses on several methodological factors as follows.

#### 6.1 Head motion threshold.

To ensure that our findings were not influenced by the potential effects of head motion [28, 29], we implemented a stricter quality control threshold, excluding participants with a mean FD exceeding 0.3 mm. Analyses of interindividual variation patterns under this criterion revealed highly consistent spatial distributions compared with the main results (mean $r = 0.99$; Fig. S15). Likewise, predictive models retrained after removing high-motion participants yielded nearly identical performance (prediction accuracy: age: $r = 0.57$, cognition: $r = 0.49$, language: $r = 0.29$, motor: $r = 0.46$, all $P_{perm} < 0.001$; contributing voxels: age: $r = -0.39$, cognition: $r = -0.43$, language: $r = -0.42$, motor: $r = -0.47$, all $P_{spin} < 0.001$; Fig. S15). These results demonstrate that our conclusions are robust against potential head motion confounds.

#### 6.2 Effect of scan duration.

Considering recent reports indicating that scan duration can substantially affect the reliability of neonatal individualized functional networks [30, 31], we tested the impact of scan length on network reconstruction. Specifically, the full times series for each infant was divided into two equal halves at the midpoint, with the first half used to generate a reference network. Then, two sampling strategies were applied to the second half to generate a series of test networks: 1) Sequential sampling with progressively increasing scan durations; and 2) Random sampling with progressively increasing scan durations. Consistent with previous studies [30, 31], network stability was quantified using the within-subject normalized mutual information (NMI) between the reference and testing networks. We found that the within-subject NMI for both group-level and individualized functional networks increased with scan duration, indicating that longer acquisition improve the stability of neonatal functional topography. Furthermore, a scan duration of 7.5 minutes achieved an acceptable level of stability comparable to existing literature [30-32] (all NMI > 0.51, Fig.S16).

#### 6.3 Sample size for brain maturity and neurodevelopmental outcome prediction.

Previous study indicated that sample size can significantly brain-behavior association studies[33, 34]. To validate the required sample size for our studies, we randomly selected participants with replacement from the full sample at logarithmically spaced sample sizes (Age: n = 315, 16 intervals: n = 25, 42, 60, 77, 94, 111, 129, 146, 163, 180, 198, 215, 232, 249, 267, 284; Neurodevelopmental outcome: n = 249, 16 intervals: n = 25, 38, 52, 65, 78, 91, 105, 118, 131, 144, 158, 171, 184, 197, 211, 224). For each sample size, 100 bootstrap replicates were used with 10-fold cross-validation, and Pearson's r between predicted and observed values quantified model performance. Our results found that a cohort of nearly 220 neonates could achieve significant predictions for both the brain maturation and all cognitive assessments, and could reach nearly 80% predictive accuracy of that from our main findings (Fig. S17). However, we only observed a convergence line for brain age prediction not for cognition predictions suggesting that a larger sample size may still valuable for the prediction of neurodevelopmental outcomes.

#### 6.4 Split-half validation.

To assess reproducibility, neonates in Subset 2 were randomly divided into two equal halves matched for scan age. The cortical distribution of functional variability was highly consistent between halves (mean r = 0.97; Fig. S18), indicating excellent internal reproducibility.

#### 6.5 Sample selection for group-level atlas generation.

To test the stability of the initial group-level functional atlases, we randomly resampled different subsets of term-born neonates (n = 10–300), following previous approaches [6, 7]. The Dice coefficient between resampled and full-sample atlases increased rapidly with sample size and stabilized above 0.8 after 50 subjects (Fig. S19), confirming the robustness of the group-level network construction.

#### 6.6 Number of functional networks.

To examine sensitivity to network resolution, we regenerated individualized functional networks with k = 9 [35, 36] and k = 17 [13, 14, 37] networks and repeated all analyses. The spatial variability patterns were highly correlated with those from the main 11-network solution (r = 0.67 for k = 9; r = 0.70 for k = 17; Fig. S20). Reconstructed predictive models also yielded consistent results (for k = 9: age r = 0.50, cognition r = 0.30, language r = 0.33, motor r = 0.35; for k = 17: age r = 0.49, cognition r = 0.32, language r = 0.32, motor r = 0.37; all $P_{perm} < 0.001$). Spatial distributions of contributing voxels were highly similar across resolutions (for k = 9: age r = -0.40, cognition r = -0.45, language r = -0.45, motor r = -0.45; for k = 17: age r = -0.25, cognition r = -0.32, language r = -0.34, motor r = -0.31; all $P_{spin} < 0.001$). These results confirm that the predictive and spatial patterns of functional topographies are robust to the choice of network resolution.

**6.7 Impact of covariate regression.**

In the primary analysis, we regressed out confounding variables from both input features and prediction targets prior to predictive modeling. Because this full-sample approach may introduce data leakage—where test samples are not entirely independent of the training process—it may influence the prediction accuracy [38]. To evaluate this effect, we repeated the prediction analyses by estimating the regression parameters exclusively within the training set and then applying these parameters to the test set.

Notably, restricting covariate regression to the training set yielded unstable parameter estimates across cross-validation folds (normalized Euclidean distance of Beta values across folds = [5.37,12.03]), likely due to the limited sample size. To address this instability, we implemented two adjustment strategies: 1) Bootstrap-based outlier exclusion: We performed bootstrap sampling of beta coefficients within the training data of each fold to derive a stable mean beta estimate. Individuals whose values deviated substantially from this stable estimate were excluded (resulting in the removal of 25% of scans). 2) Robust regression: All individuals were retained in the prediction analysis, but a robust regression [39] was employed in place of the original regression model to control for covariates. Under both strategies, the predictive performance of all models remained statistically significant (Bootstrap exclusion: Age: r = 0.50, Cognition: r = 0.27, Language: r = 0.19, Motor: r = 0.22, Q-chat: r = 0.19, Preterm classification: Acc = 0.83, all $P_{\text{perm}} < 0.01$; Robust regression: Age: r = 0.47, Cognition: r = 0.16; Language: r = 0.11; Motor: r = 0.15; Q-chat = 0.12; Preterm classification: Acc = 0.77, all $P_{\text{perm}} < 0.03$, Fig.S21).

**6.8 Matched-sample validation of brain age gap analysis.**

To address potential biases arising from uneven sample, we selected term neonates that matched 52 preterm infants based on scan age, sex, and mean FD and re-estimated the BAG of matched term-born and preterm individuals, we observed highly consistent results at both overall brain and network level. Preterm infants exhibited significantly greater BAGs than matched term-born controls at both the whole-brain and network levels, confirming the robustness of our findings (Fig. S22).

### Supplementary figures

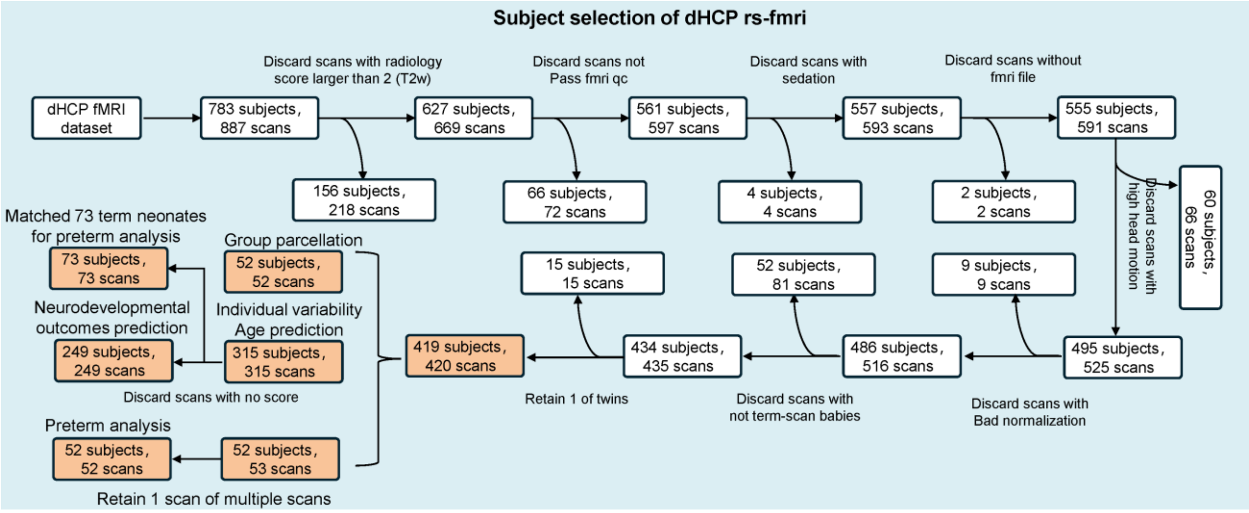


**Fig. S1 Flowchart illustrating the inclusion/exclusion criteria for MR images.** Among the initial 783 participants, 156 were excluded because of low radiological scores (ranging from 3 to 5). A total of 66 participants were excluded because of failure to pass the fMRI quality control. 4 participants and 2 participants were excluded because of sedation and a lack of fMRI files, respectively. 60 participants were excluded because of excessive head motion (head-motion displacement >5 mm, rotation >5°, or mean framewise displacement (mean FD > 0.5 mm)). 9 participants were excluded because of poor registration quality. 52 participants were excluded because they were born prematurely and did not complete a full-term scan. 15 participants were excluded because they were twins. After rigorous quality control, a total of 419 participants remained. Among these 419 participants, 367 were term-born healthy neonates, and 52 were preterm-born infants at term-equivalent age. From these 367 term-born healthy neonates, 52 neonates were used to create the group atlas, while the remaining 315 participants were employed for studying individual variability and predicting brain maturity. Given that not all participants had cognition, language, and motor scores available at 18 months, we ultimately selected 249 participants for the prediction of language and motor scores at 18 months. To examine aberrant brain maturation in preterm infants, we used data from 52 preterm infants and selected 73 term-born neonates matched for scan age, sex, and head motion to serve as controls.

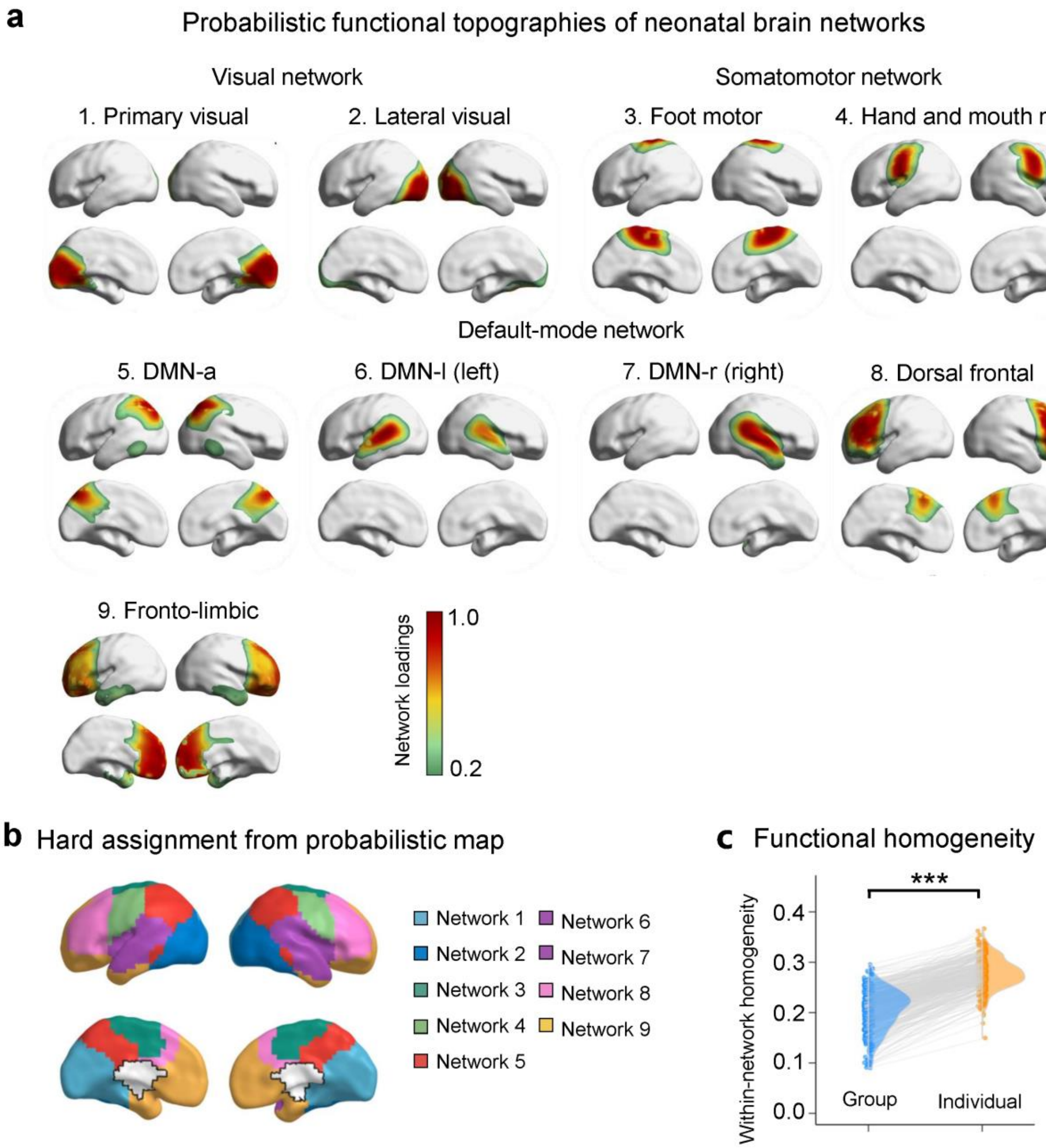


**Fig. S2 Generation of neonatal functional networks with network number k = 9.** a, The probabilistic atlas with 9-network decomposition includes primary visual (Network 1), lateral visual (Network 2), foot motor (Network 3), hand and mouth motor (Network 4), posterior DMN (Network 5), left and right lateral DMN (Networks 6–7), dorsal frontal (Network 8), and fronto-limbic (Network 9) networks. While sensorimotor networks exhibit clear spatial boundaries, higher-order association networks remain spatially diffuse and fragmented. Probabilistic loading maps indicate the degree to which each voxel is associated with a given network (red = high loading, green = low loading). **b,** Hard parcellation maps obtained by assigning each voxel to the network with the highest probability loading. **c,** Functional homogeneity was significantly greater in individualized networks than in group-level networks, confirming the effectiveness of personalized network delineation. ***, $P < 0.001$. Source data are provided as a Source Data file.

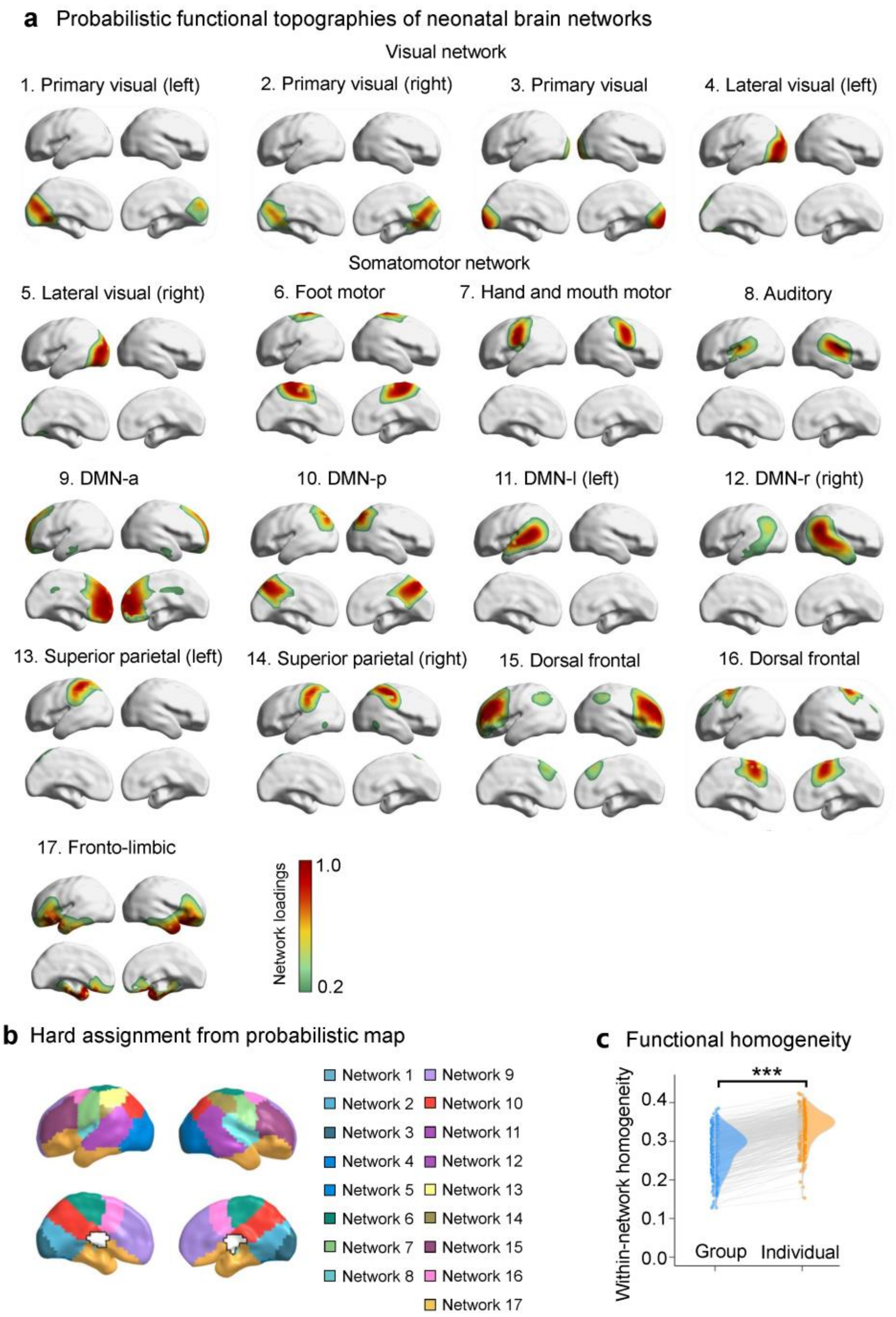


**Fig. S3 Generating neonatal functional networks with network number k = 17.** **a,** The group-level probabilistic atlas derived from k = 17 decomposition identified well-differentiated sensorimotor systems, including bilateral primary visual (Networks 1–3), lateral visual (Networks 4–5), foot motor (Network 6), hand and mouth motor (Network 7), and auditory cortex (Network 8) networks, as well as distributed association networks, including anterior DMN (DMN-a, Network 9), posterior DMN (DMN-p, Network 10), lateral DMN (DMN-l and DMN-r, Networks 11–12), bilateral superior parietal (Networks 13–14), dorsal frontal (Networks 15–16), and fronto-limbic region (Network 17) networks. Probabilistic maps depict the voxel-wise network loading strength (red = high, green = low). **b,** Hard parcellation maps were created by assigning each voxel to the network with the highest probability loading. **c,** Functional homogeneity was significantly greater in individualized maps than in group-level parcellations, supporting the advantage of personalized network delineation. ***, $P < 0.001$. Source data are provided as a Source Data file.

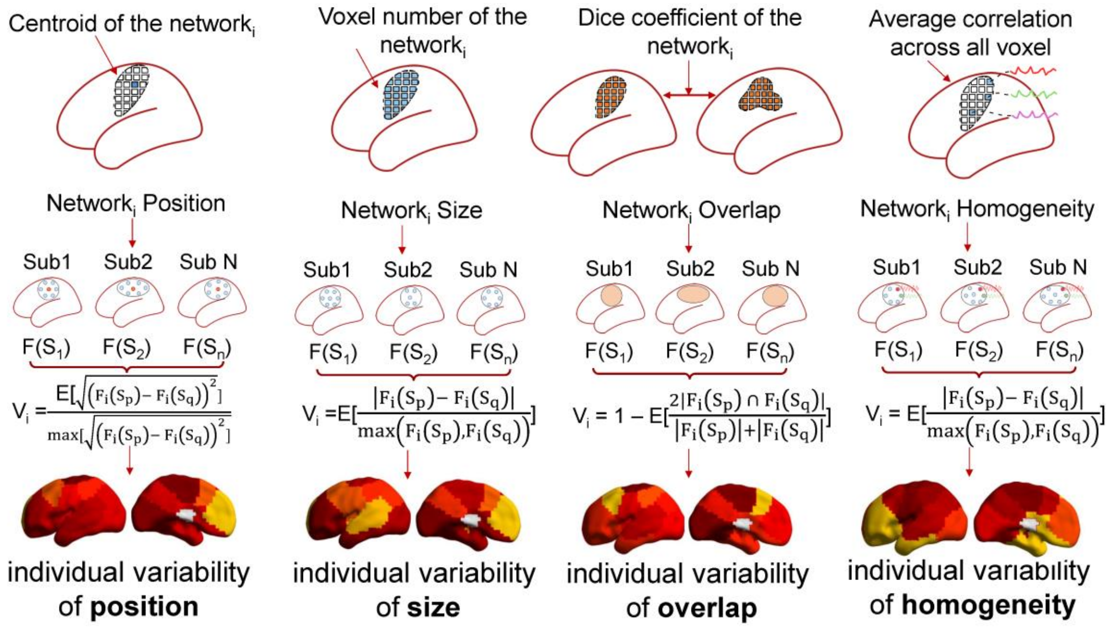


**Fig. S4 Definition of network-level topographic variability across individuals**. (1) Position variability was defined as the average Euclidean distance between network centroids (centres of mass) across participants, normalized by the maximum average Euclidean distance. (2) Size variability was quantified as the standard deviation of the voxel counts per network across participants, normalized by the maximum size. (3) Overlap variability was defined as one minus the average Dice coefficient between network pairs, capturing spatial correspondence variability across individuals. (4) Homogeneity variability was measured as the standard deviation of within-network functional homogeneity, normalized by the maximum homogeneity. Group-level comparisons between primary and association networks were evaluated using 10,000 spin permutations. For methodological details, see (ref.[40]).

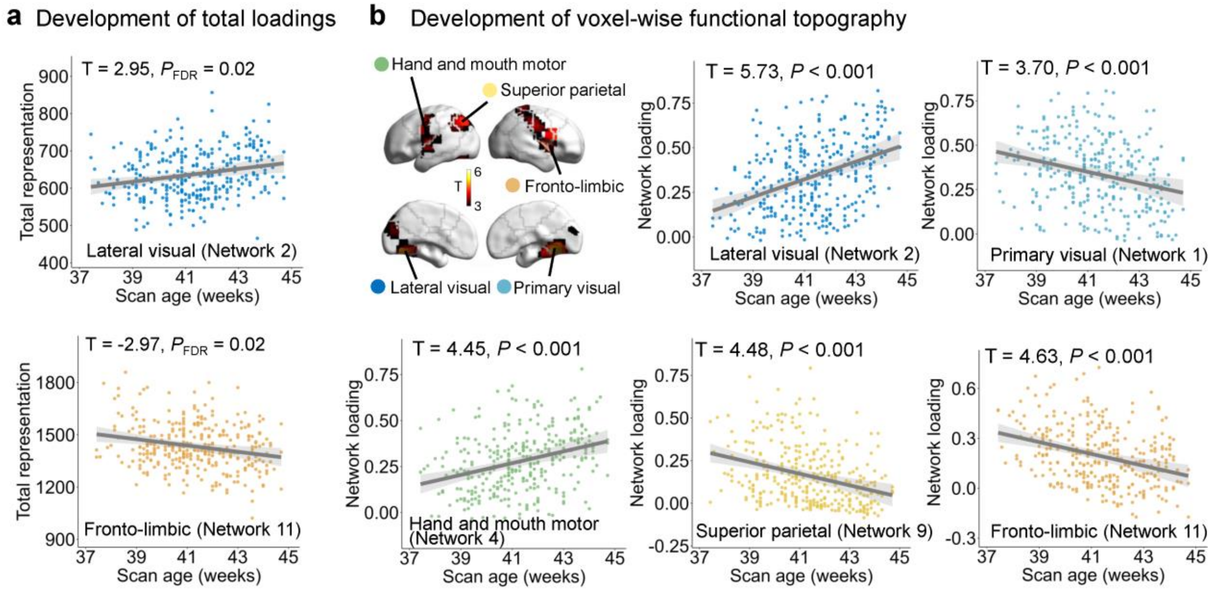


**Fig. S5 Developmental changes in both global and voxel-wise network organization using generalized linear models (GLMs). a,** Developmental trajectories of total network representation across scan ages. The total representation was quantified as the sum of the voxel-wise probabilistic loadings for each network. **b,** Developmental trajectories of the voxel-wise functional topography. Analyses were conducted at the voxel level ($P < 0.001$) and corrected at the cluster level ($P < 0.05$) using Gaussian random field theory. Shaded areas denote 95% confidence intervals. Source data are provided as a Source Data file.

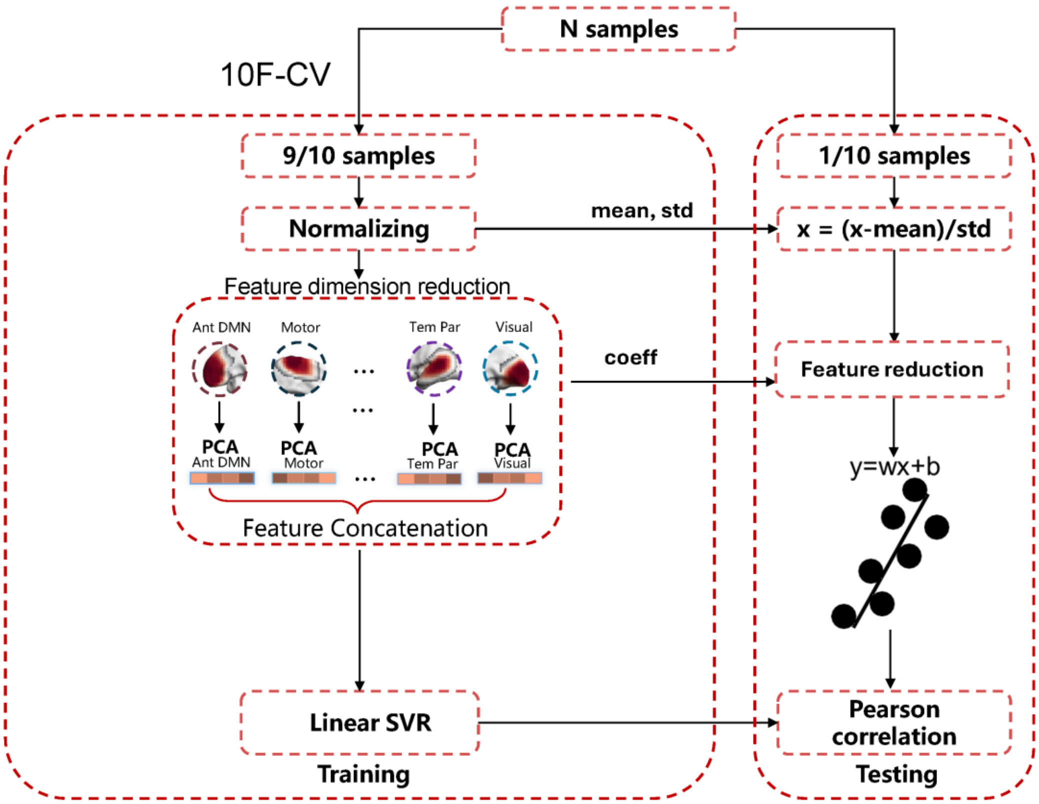


**Fig. S6 Flowchart of the Ti-PCA prediction framework with 10F-CV.** The Ti-PCA framework was developed to assess whether individualized functional topography can predict brain maturity and neurodevelopmental outcomes. Voxel-wise functional topographies were first reduced using network-wise PCA, preserving within-network topographic independence while minimizing dimensionality. The resulting low-dimensional features across all the networks were concatenated and used to train an SVR model. The models were trained and tested with a 10F-CV approach. Prediction performance was quantified by the Pearson correlation (r) between the predicted and actual values in the testing set.

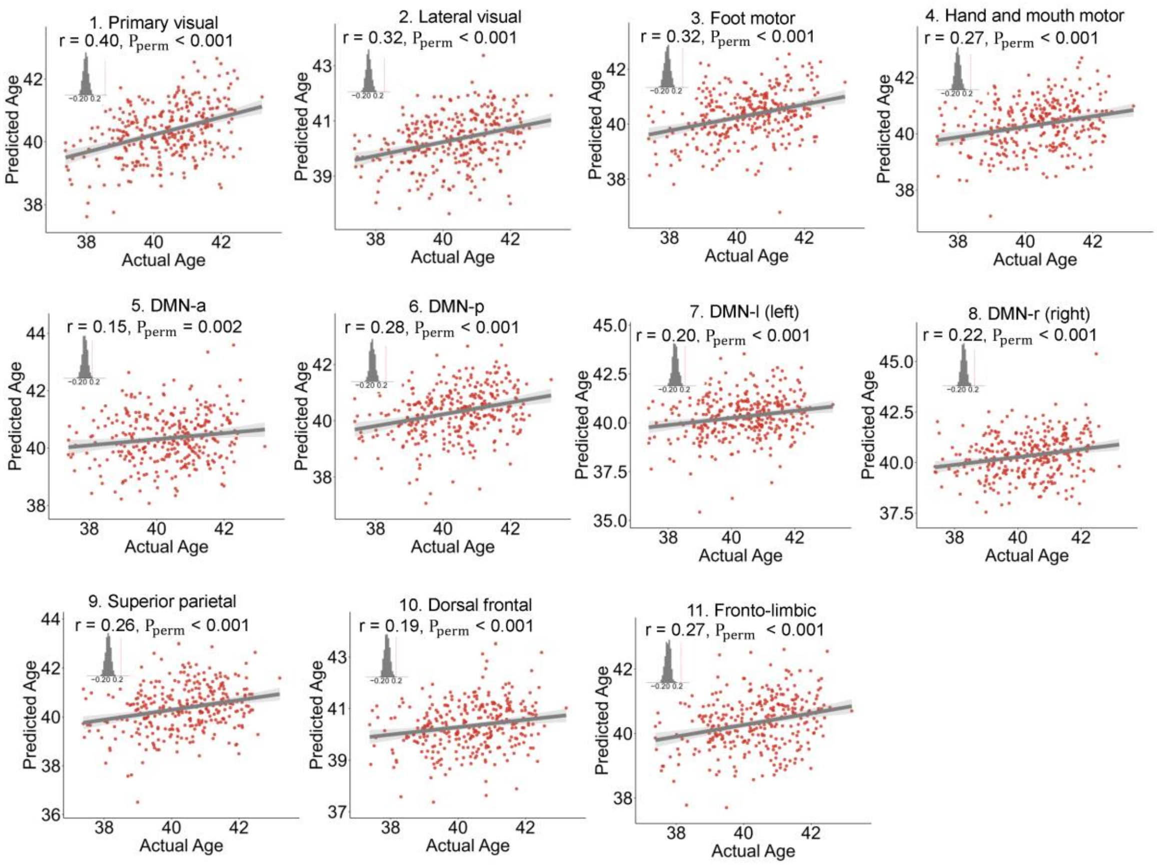


**Fig. S7 The prediction performance of each functional network separately for individual brain maturity.** All 11 networks showed significant predictive performance, with primary networks yielding the highest accuracy. Each data point represents an individual's predicted score from a model trained on held-out participants. Insets show null distributions from 1,000 permutation tests. Shaded areas denote 95% confidence intervals. Source data are provided as a Source Data file.

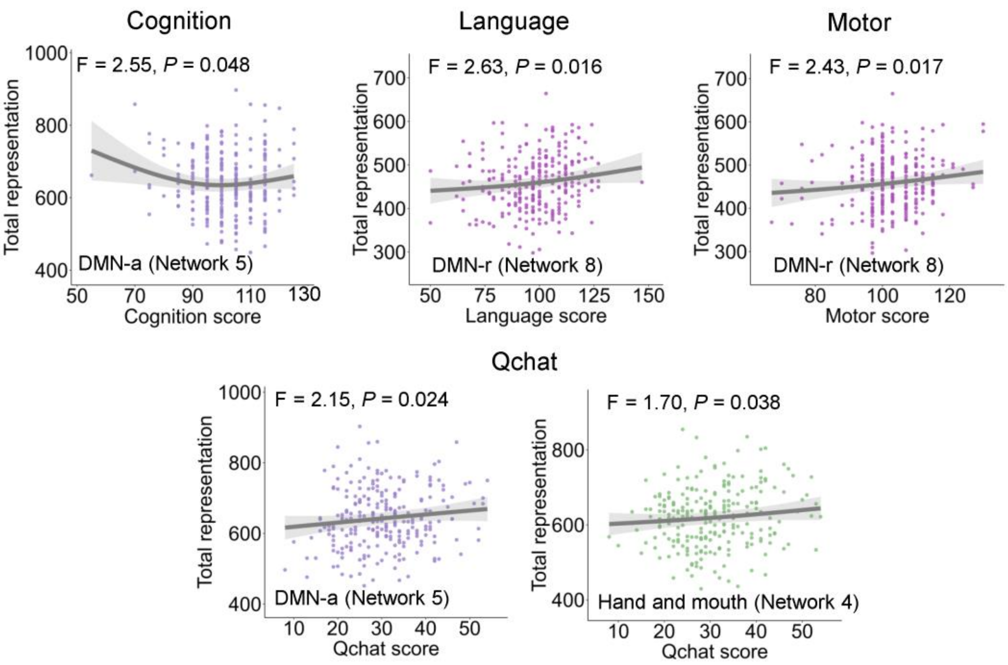


**Fig. S8 Associations between network representation and 18-month neurodevelopmental outcomes (uncorrected *P* values).** Shaded areas denote 95% confidence intervals. Source data are provided as a Source Data file.

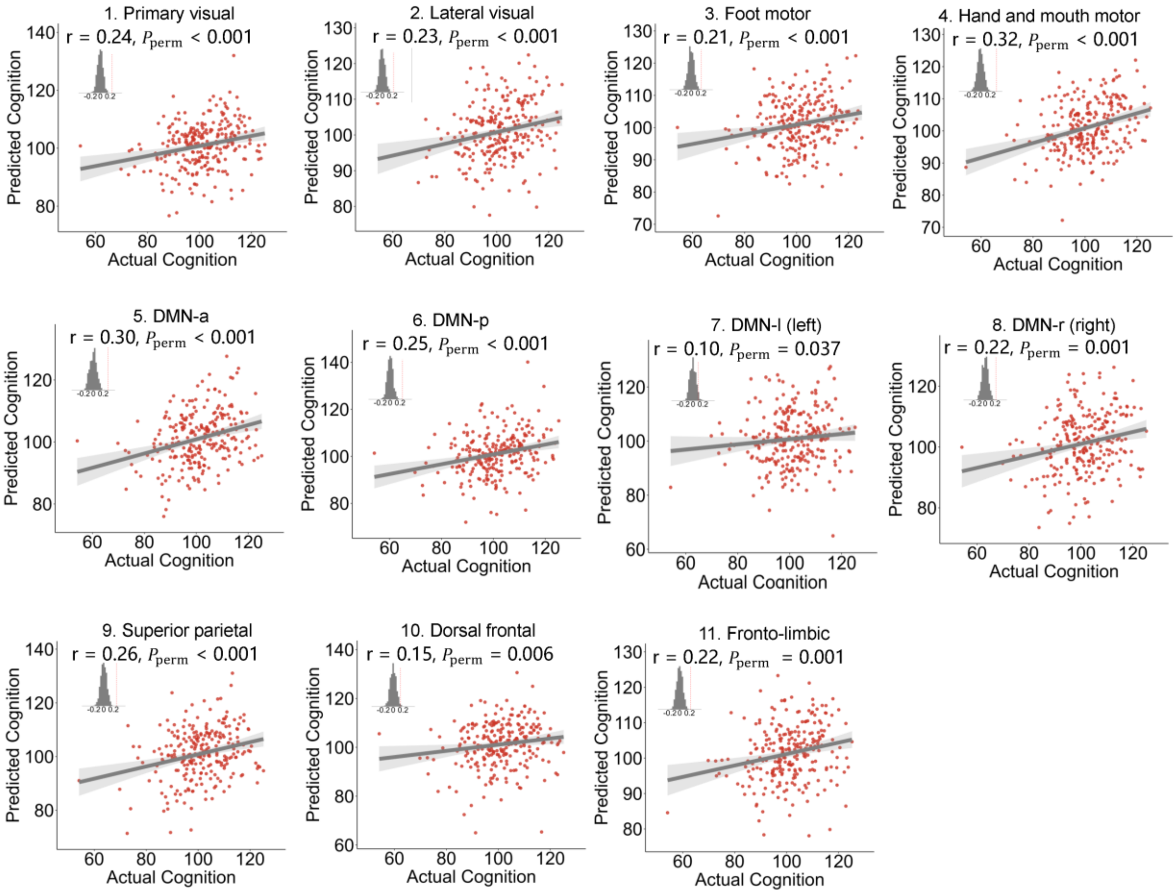


**Fig. S9 The prediction performance of each functional network for individual cognition scores at 18 months.** All 11 networks showed significant predictive performance, with primary networks yielding the highest accuracy. Each data point represents an individual's predicted score from a model trained on held-out participants. Insets show null distributions from 1,000 permutation tests. Shaded areas denote 95% confidence intervals. Source data are provided as a Source Data file.

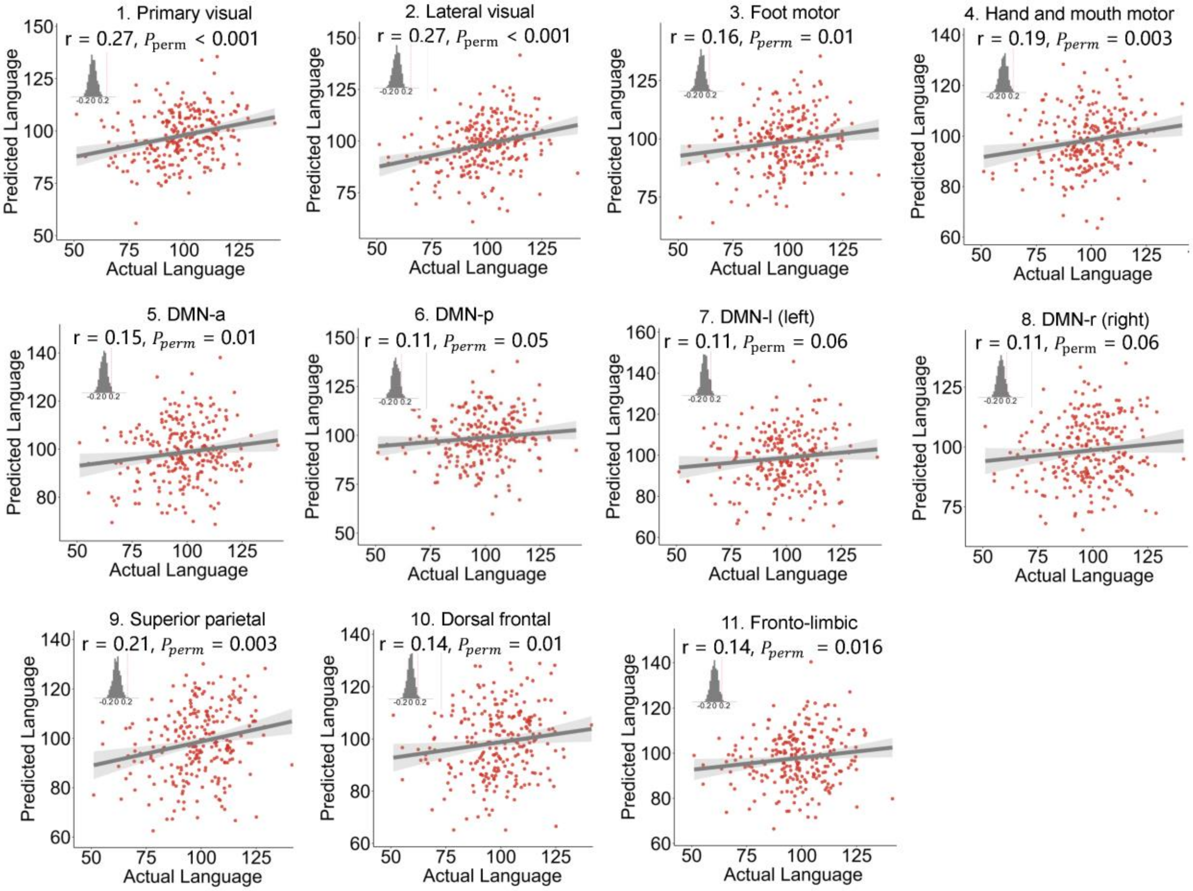


**Fig. S10 The prediction performance of each functional network separately for individual language scores at 18 months.** Most networks showed significant predictive performance, with primary networks yielding the highest accuracy. Each data point represents an individual's predicted score from a model trained on held-out participants. Insets show null distributions from 1,000 permutation tests. Shaded areas denote 95% confidence intervals. Source data are provided as a Source Data file.

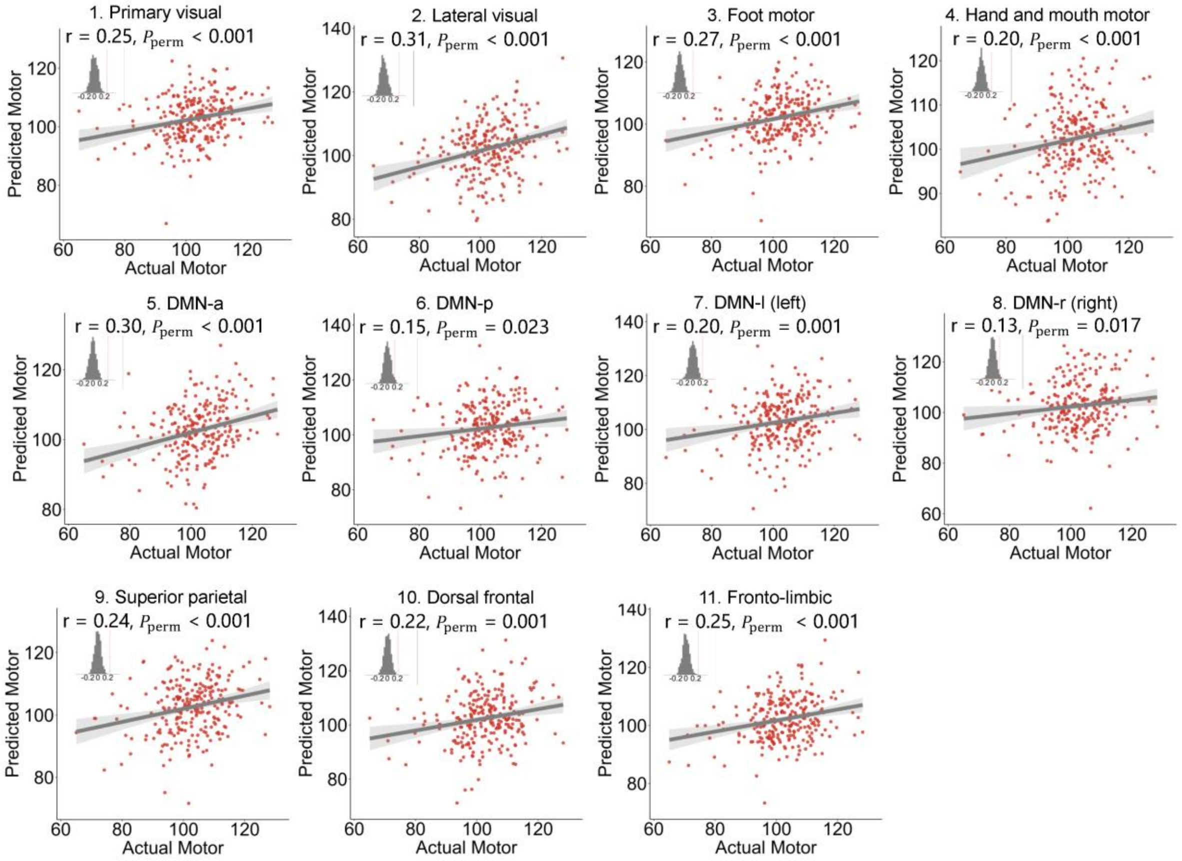


**Fig. S11 The prediction performance of each functional network separately for individual motor scores at 18 months.** All 11 networks showed significant predictive performance, with primary networks yielding the highest accuracy. Each data point represents an individual's predicted score from a model trained on held-out participants. Insets show null distributions from 1,000 permutation tests. Shaded areas denote 95% confidence intervals. Source data are provided as a Source Data file.

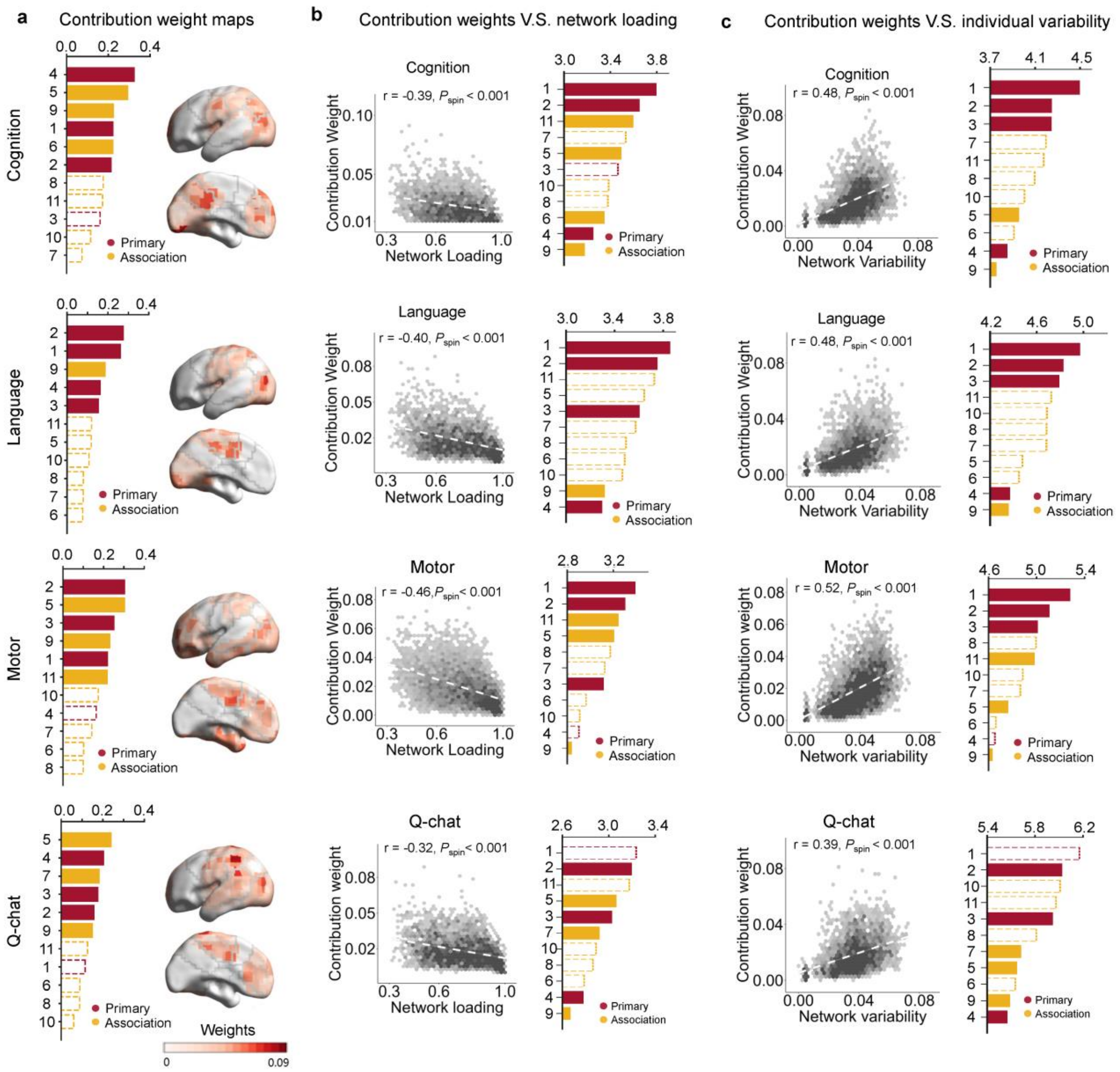


**Fig. S12 Prediction contribution weights are primarily concentrated in regions with low network loadings and high interindividual variability. a,** Contribution weight maps for the prediction of cognitive, language, and motor outcomes before partial regression analyses. The bars indicate network-level weights (red: primary; yellow: association). Brain maps depict voxel-wise predictive weights. **b,** Voxel-wise contribution weights were negatively correlated with network loading. **c,** Voxel-wise contribution weights were positively correlated with interindividual network variability. Dashed outlines denote networks that were not classified as high-contributing networks (z-score < 0). Shaded areas denote 95% confidence intervals. Source data are provided as a Source Data file.

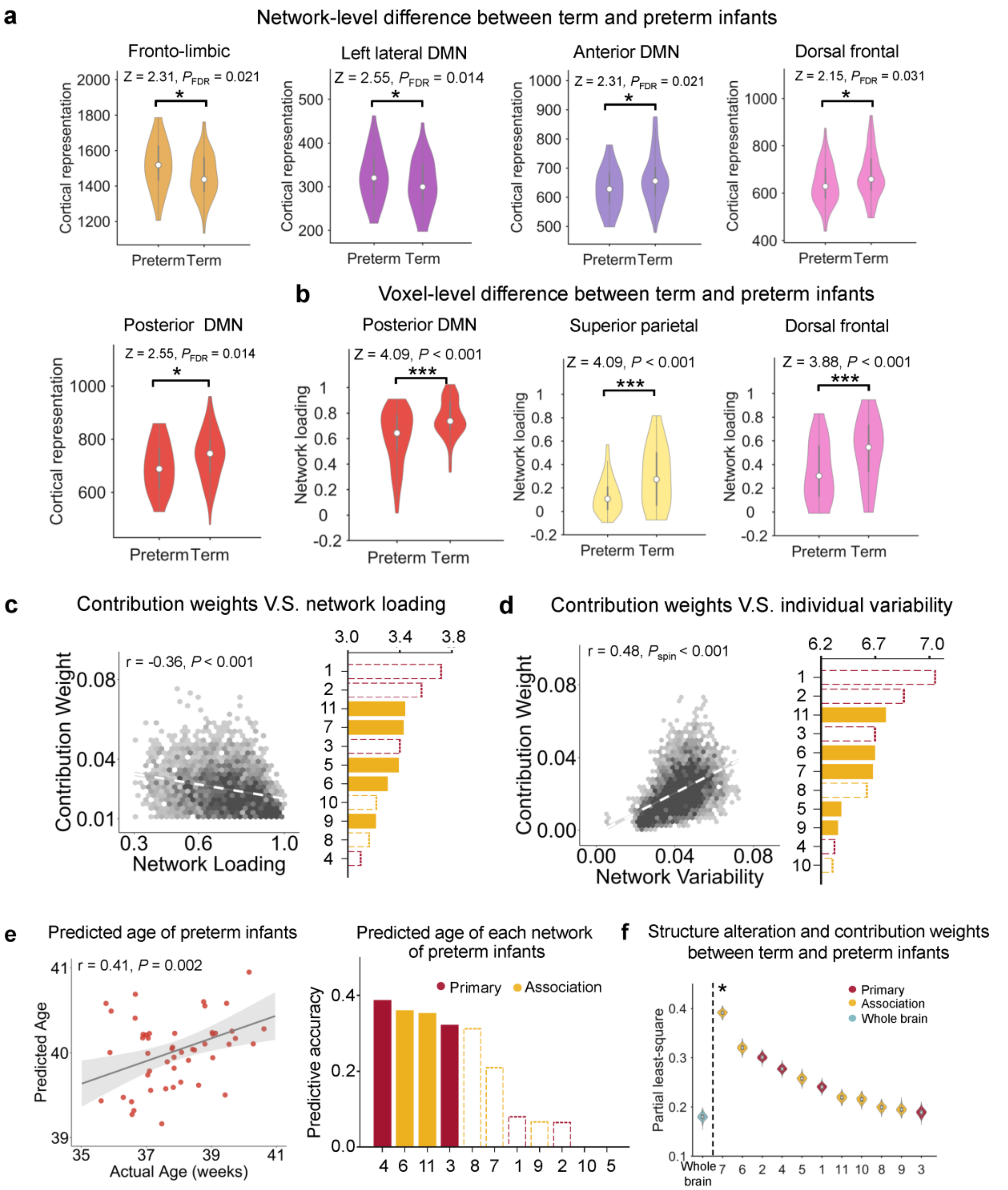


**Fig. S13 Results of the estimation of abnormal functional topography in preterm infants. a,** Network-level differences in total network representation between term and preterm infants. **b,** Voxel-wise topographic differences between term and preterm infants. **c,** Contribution weights derived from the TiPCA classification model were negatively correlated with network loading. **d,** Contribution weights were positively correlated with interindividual topographic variability. Bar plots summarize these correlations across networks. **e,** Brain age prediction in preterm infants using models trained on term-born neonates for whole-brain and network-level topographies. **f,** PLS analysis revealed the association between structural abnormalities and functional classification weights. Dashed outlines denote networks that were not classified as high-contributing networks (*z*-score < 0). *, *P* < 0.05; ***, *P* < 0.001. Shaded areas denote 95% confidence intervals. Source data are provided as a Source Data file.

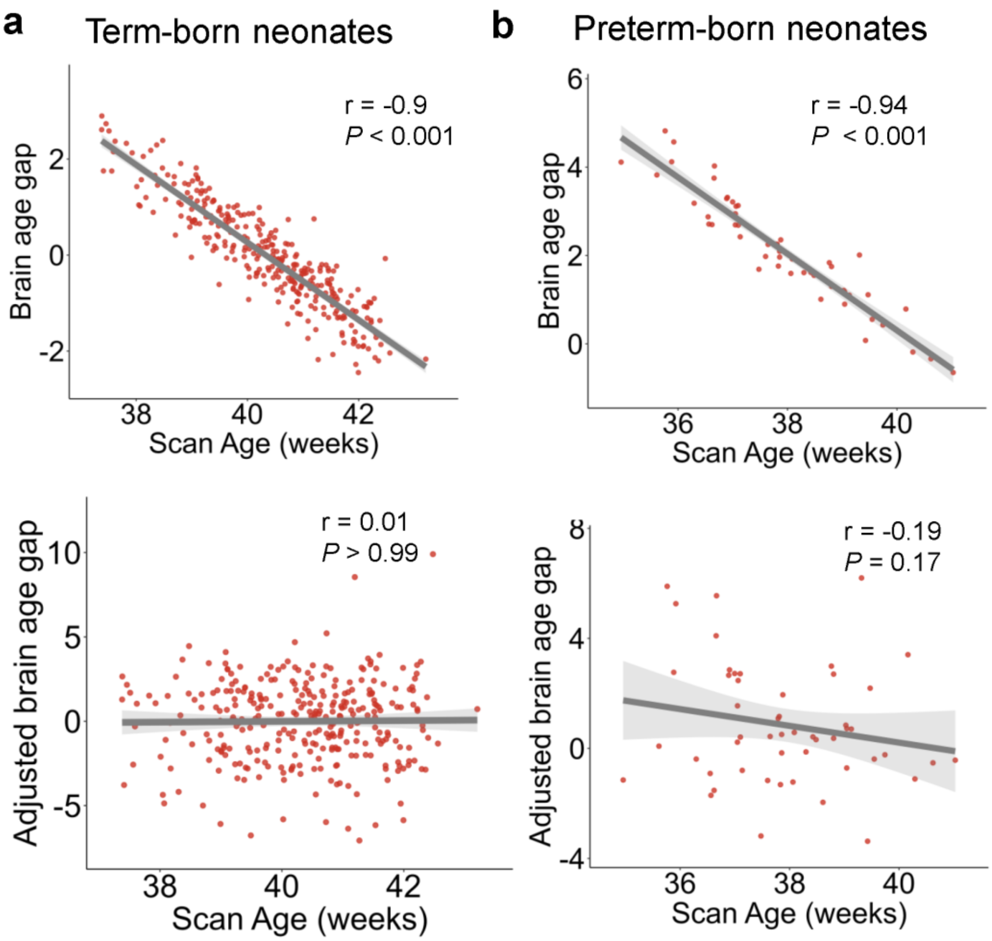


**Fig. S14 Adjustment of the brain age gap (BAG) to correct for regression-to-mean bias [41]. a,** Predicted age was corrected by the slope and intercept of fitting in term-born neonates. Before correction, the predicted BAG was significantly associated with scan age (upper panel); After correction, the predicted BAG was no longer significantly associated with scan age (bottom panel). **b,** For the preterm-born neonates, we directly employed the correction coefficients obtained in term-born samples. Before correction, the predicted BAG was significantly associated with scan age (upper panel). After correction, the adjusted BAG was also no longer significantly associated with scan age (bottom panel). This correction enables unbiased comparisons of BAG between term- and preterm-born groups. Shaded areas denote 95% confidence intervals. Source data are provided as a Source Data file.

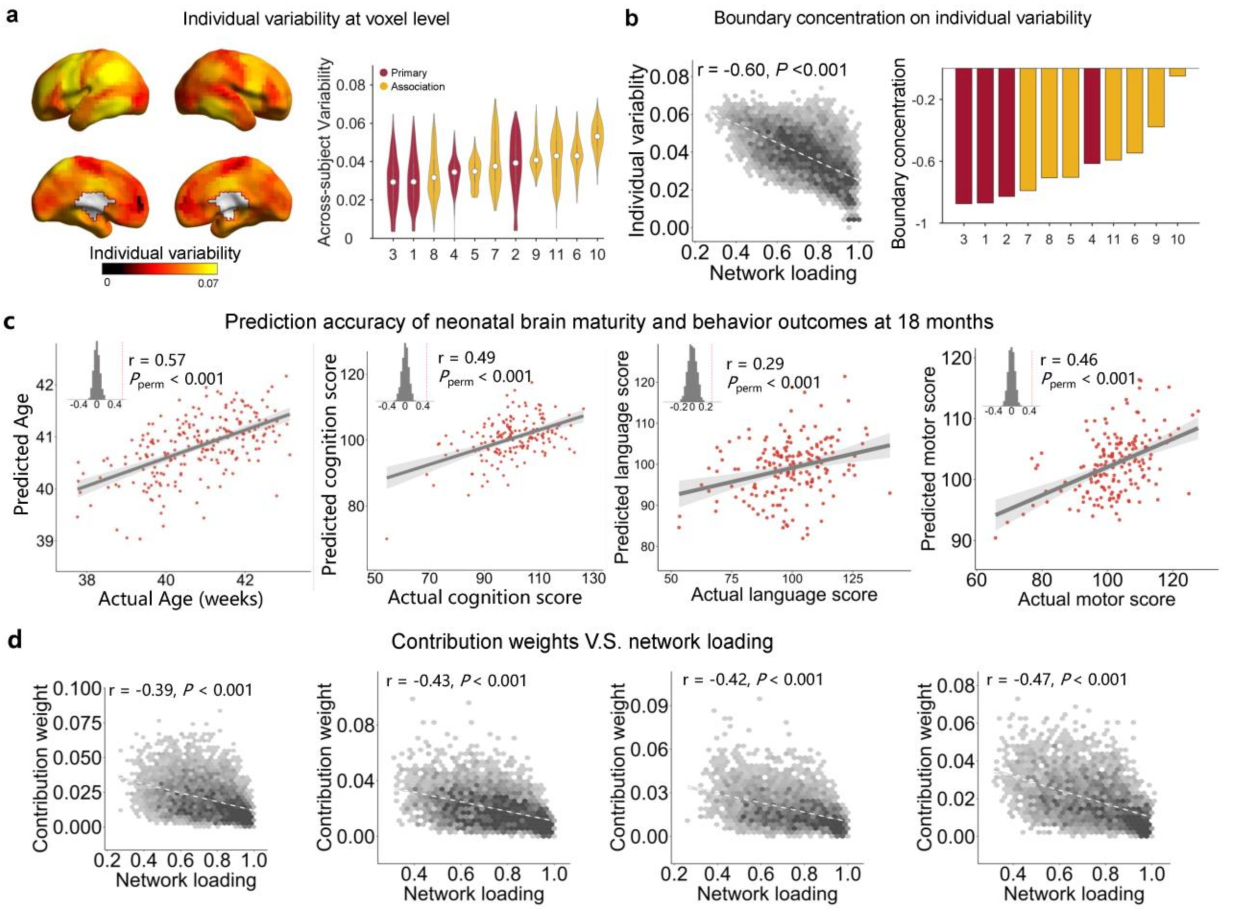


**Fig. S15 Individual variability and predictive relevance of neonatal functional topography in selected participants under a stricter FD threshold (mean FD < 0.3 mm). a,** Voxel-wise variability was greater in association regions than in primary regions (r = 0.9945 with main result, $P < 0.001$), confirming greater interindividual differences in association cortices. **b,** Boundary concentration analysis revealed that regions with the greatest interindividual variability were preferentially near network borders. **c,** Complex patterns of functional topography could be used to predict brain maturity and cognitive, language, and motor abilities at 18 months in unseen data. **d,** Contribution weights were negatively correlated with network loading. Shaded areas denote 95% confidence intervals. Source data are provided as a Source Data file.

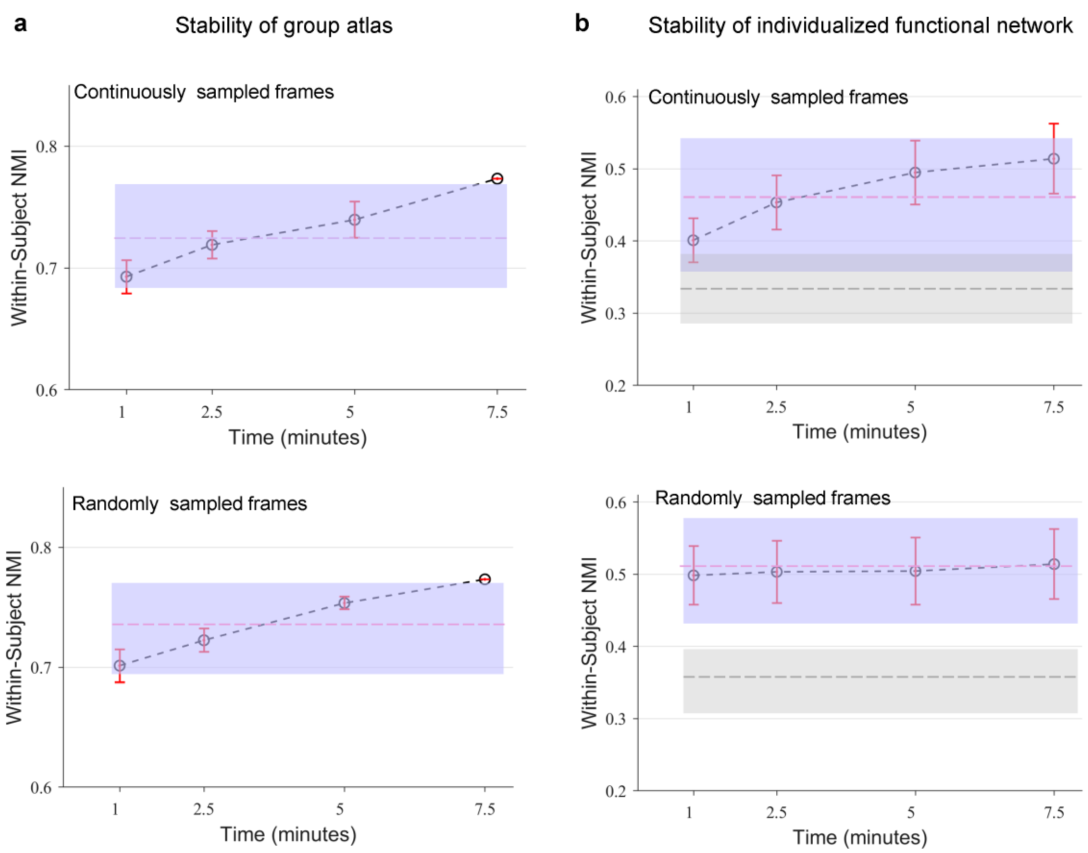


**Fig. S16 Required scan duration for generating stable group-level and individualized functional networks. a,** Stability (within-individual NMI) of group-level functional networks generated from different scan durations using continuously sampled frames (top) and randomly sampled frames (bottom). **b,** Stability (within-individual NMI) of individualized functional networks generated from different scan durations using continuously sampled frames (top) and randomly sampled frames (bottom). Of note, the purple boxes display the 5th to 95th percentiles of within-individual NMI values across all individuals and all samplings, while the gray boxes display the same for between-individual NMI (the average NMIs are displayed as dashed lines in each colour). Source data are provided as a Source Data file.

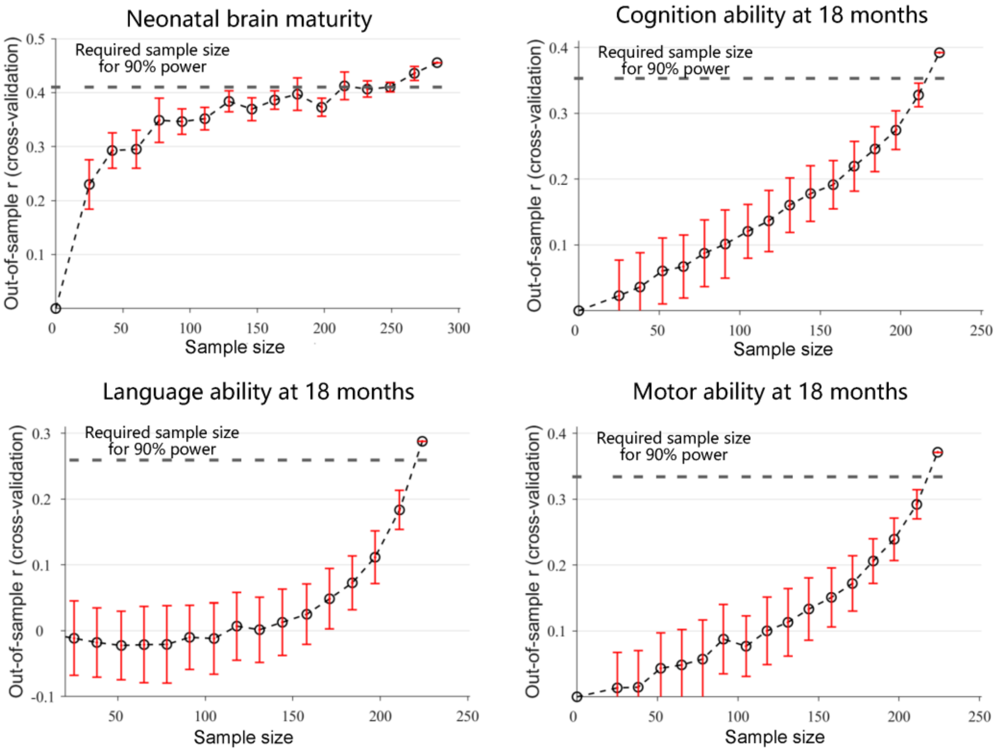


**Fig. S17 Required sample size for the Ti-PCA-based prediction of neonatal brain and cognitive outcomes.** In a cohort of nearly 220 neonates, significant predictions were achieved for both brain maturation and all cognitive assessments, reaching nearly 90% predictive accuracy of that from our main findings. However, we observed only a convergence line for brain age prediction and not for cognition prediction, suggesting that a larger sample size may still be valuable for the prediction of neurodevelopmental outcomes. Source data are provided as a Source Data file.

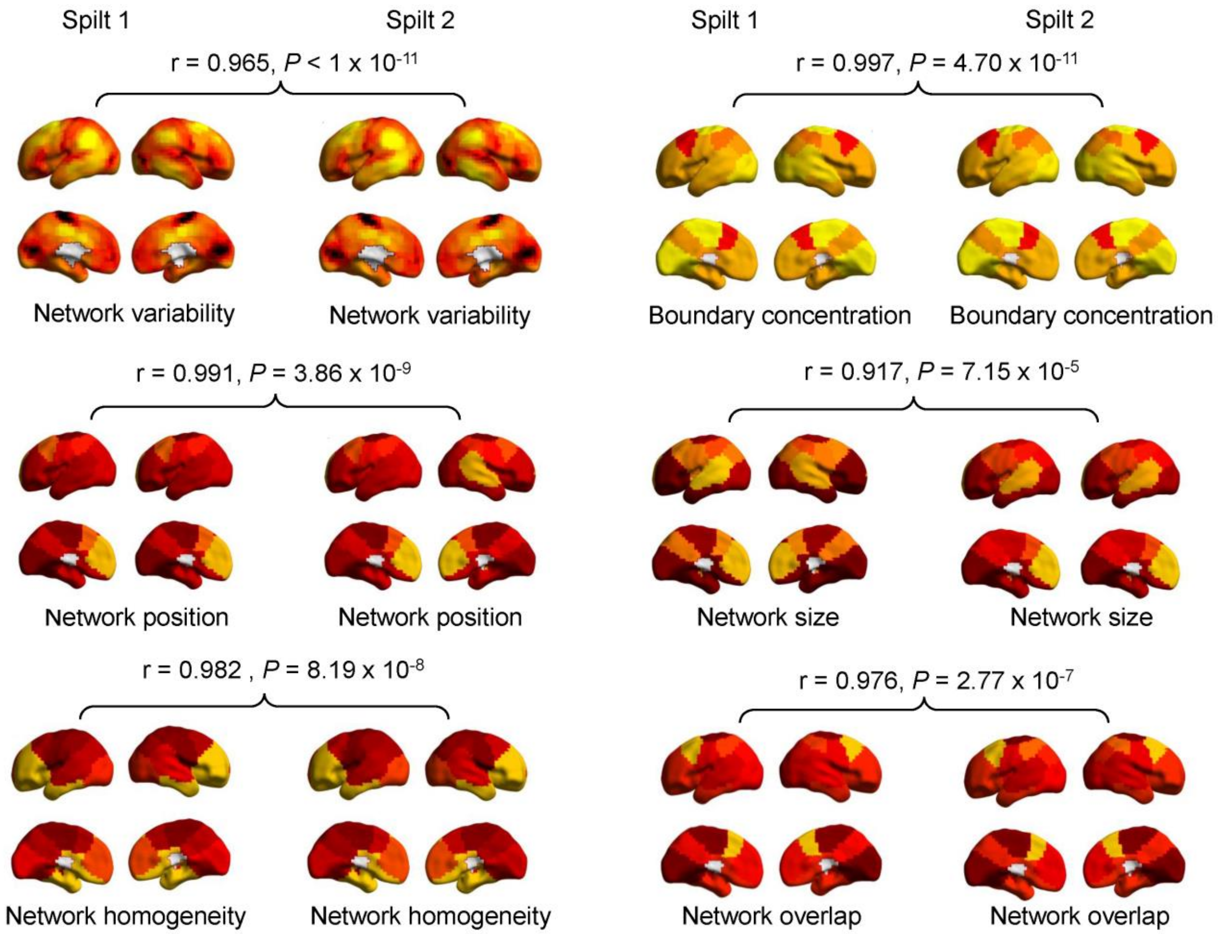


**Fig. S18 Reproducibility of functional topographic variability in split-half analysis.** To evaluate the reproducibility of functional topographic variability, 315 term-born neonates (Subset 2) were randomly split into two groups matched by scan age. All the voxel-wise and network-level variability metrics—including network variability, boundary concentration, and four topographic properties (position, size, overlap, and homogeneity)—were independently computed for each half. Pairwise spatial correlations were used to assess reliability across pairs. Source data are provided as a Source Data file.

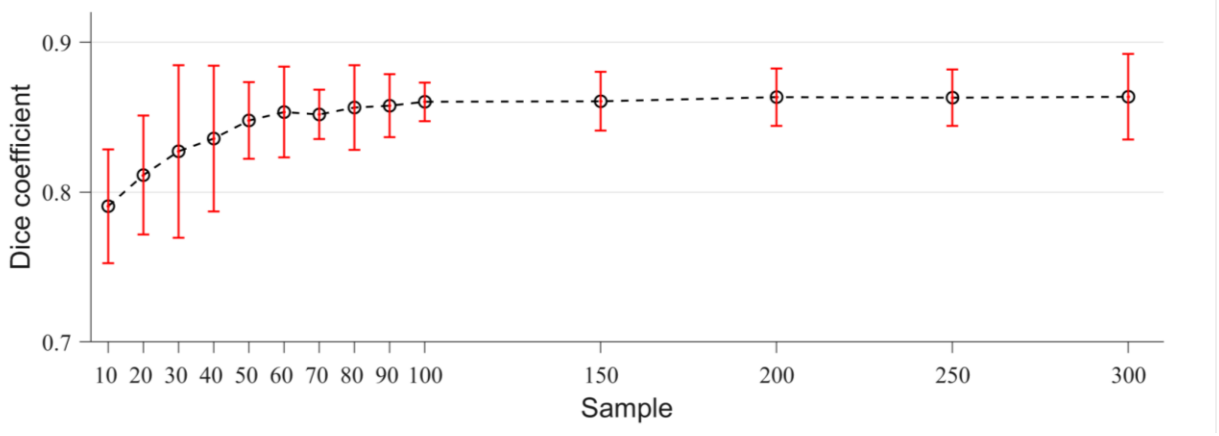


**Fig. S19 Stability of group-level functional networks under varying sample sizes.** To assess the impact of sample size on group-level functional network generation, we randomly selected 10 to 300 term-born neonates and regenerated group-level atlases. For each sample size, the procedure was repeated 10 times, and the Dice coefficient was computed between the subsample-derived atlas and the full-sample reference. The error bars indicate the standard deviation across repetitions. The Dice coefficient increased with sample size and stabilized above 0.85 after approximately 50 participants, demonstrating the robustness of group-level network construction. Source data are provided as a Source Data file.

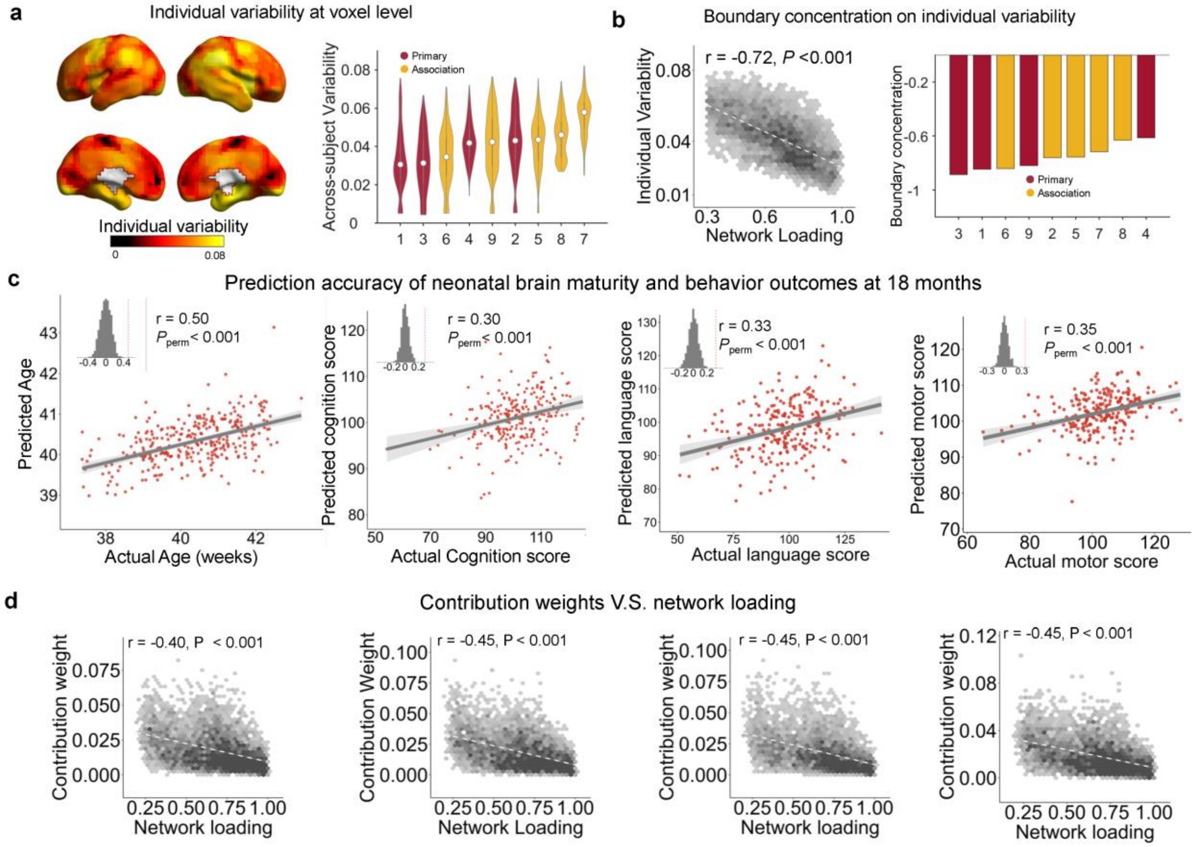


**Fig. S20-I Individual variability and predictive relevance of neonatal functional topography with 9-network decomposition. a,** Voxel-wise variability was greater in association regions than in primary regions (r = 0.67 with main result, $P < 0.001$), confirming greater interindividual differences in association cortices. **b,** Boundary concentration analysis revealed that regions with the greatest interindividual variability were preferentially near network borders. **c,** The complex pattern of functional topography could be used to predict brain maturity and 18-month cognitive, language, and motor outcomes in unseen data. **d,** Contribution weights were negatively correlated with network loading, indicating that regions near network boundaries contributed most to the prediction of brain maturity and neurodevelopmental outcomes. Shaded areas denote 95% confidence intervals. Source data are provided as a Source Data file.

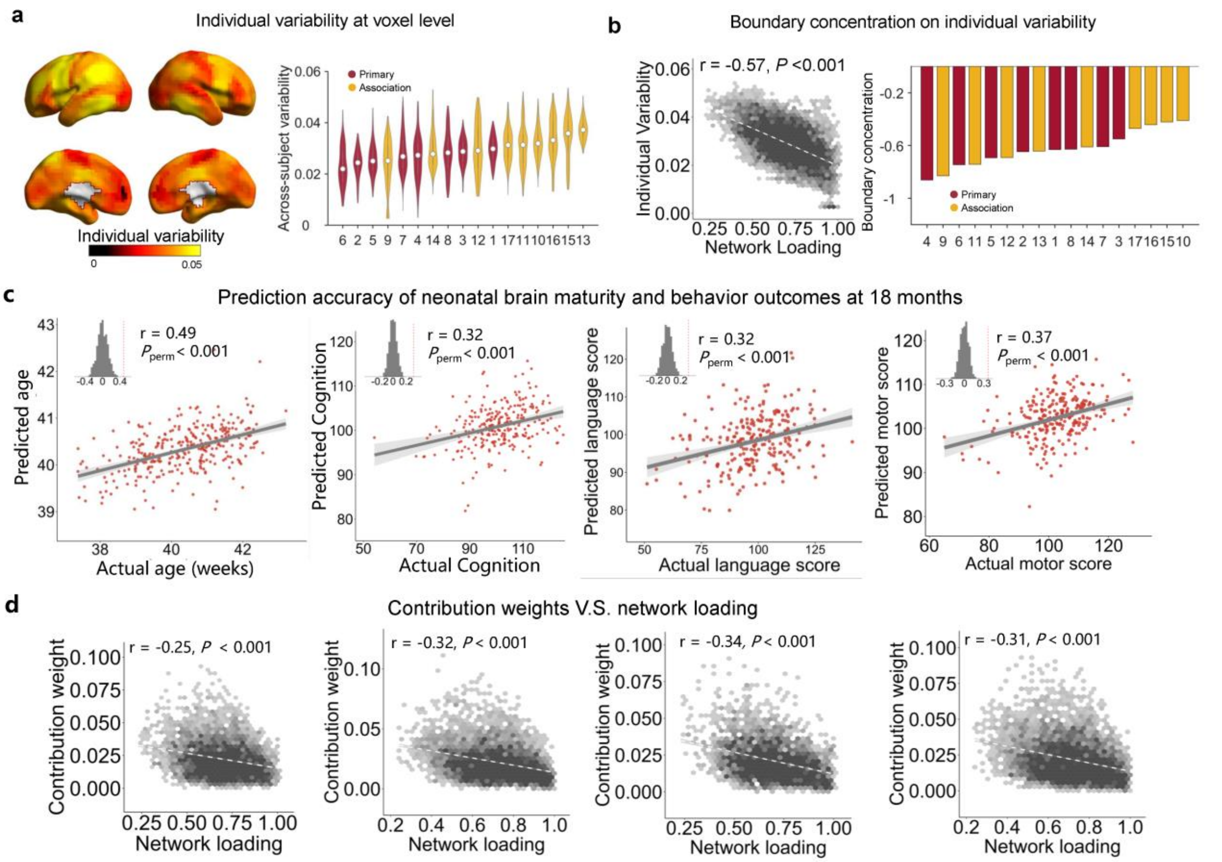


**Fig. S20-II Individual variability and predictive relevance of neonatal functional topography with 17-network decomposition. a,** Voxel-wise variability was greater in association regions than in primary regions (r = 0.70 with main result, *P* < 0.001), confirming greater interindividual differences in association cortices. **b,** Boundary concentration analysis revealed that regions with the greatest interindividual variability were preferentially near network borders. **c,** The complex pattern of functional topography could be used to predict brain maturity and cognitive, language, and motor abilities at 18 months in unseen data. **d,** Contribution weights were negatively correlated with network loading, indicating that regions near network boundaries contributed most to the prediction of brain maturity and neurodevelopmental outcomes. Shaded areas denote 95% confidence intervals. Source data are provided as a Source Data file.

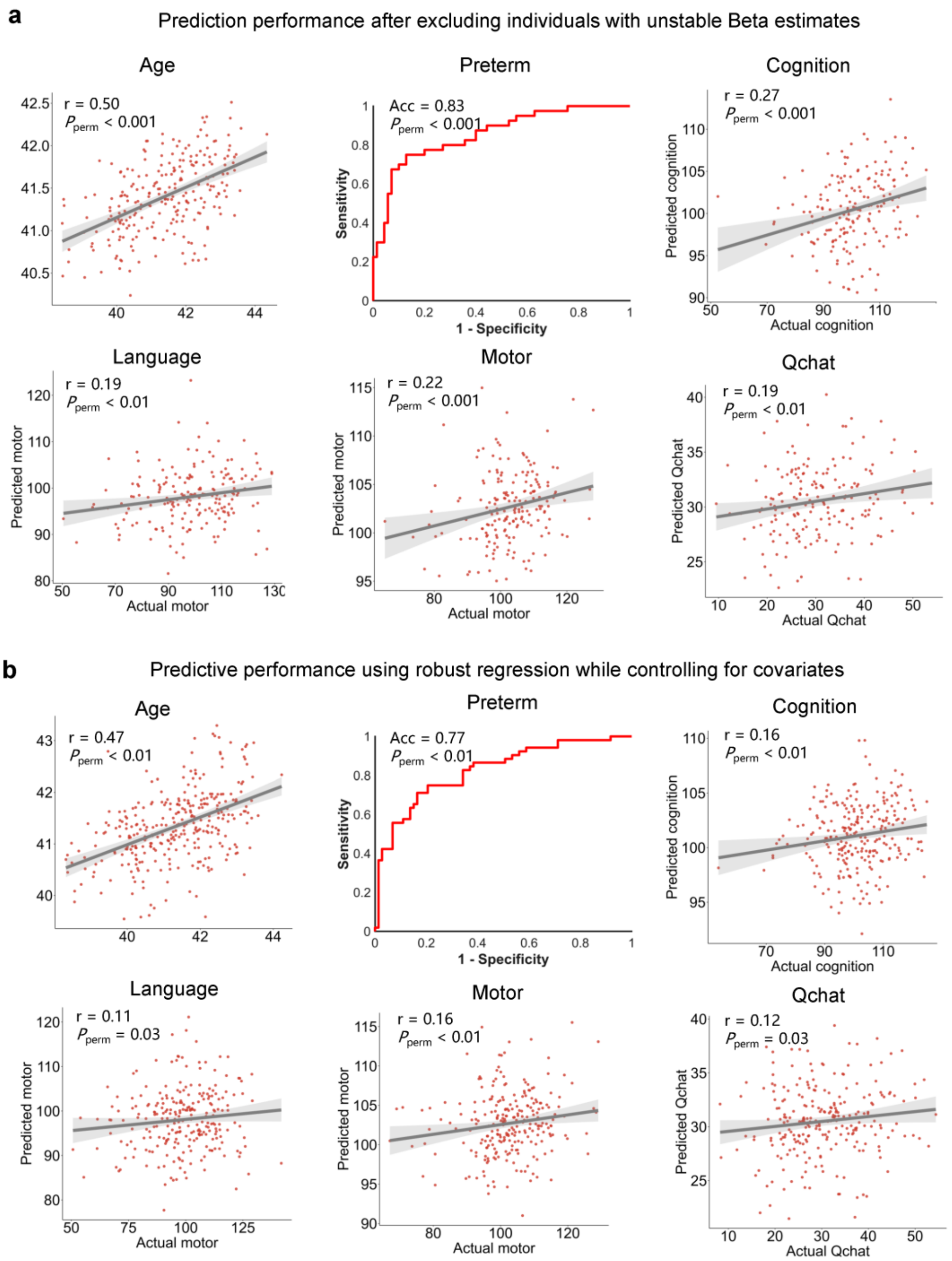


**Fig. S21 Prediction and classification performance when confound regression was performed only in the training set. a,** Prediction performance by excluding individuals who deviated largely from a stable Beta obtained by bootstrap sampling. **b,** Prediction performance by employing a robust regression in all individuals. Each dot represents one participant. Solid lines indicate linear fits and shaded areas represent 95% confidence intervals. Shaded areas denote 95% confidence intervals. Source data are provided as a Source Data file.

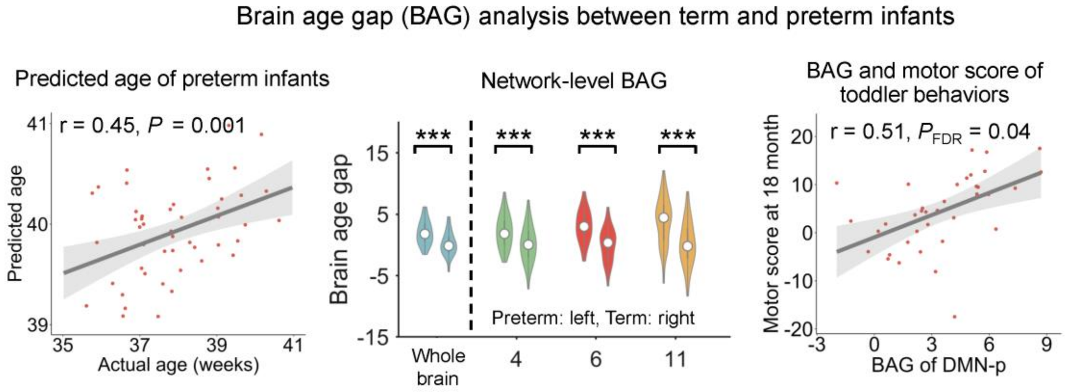


**Fig. S22 BAG analysis of preterm infants based on matched term-born neonates.** To control for sample heterogeneity, 52 preterm infants were matched to 52 term-born controls on the basis of scan age, sex, and head motion. A model trained on term-born neonates predicted brain age in preterm infants with significant accuracy (r = 0.45, *P* = 0.001, left). BAG was significantly elevated in preterm infants at both the whole-brain and network levels, with the greatest effects observed in the posterior default mode network (DMN-p), fronto-limbic network, and hand/mouth motor network (middle). The BAG within the DMN-p was positively associated with 18-month motor outcomes (r = 0.51; $P_{FDR}$ = 0.001; right), indicating a link between accelerated functional maturation and later behavioural development. Shaded areas denote 95% confidence intervals. Source data are provided as a Source Data file.

**Table S1 Demographic and clinical characteristics of term and preterm infants.**

| | Term | Preterm | Test statistic | P |
|---|---|---|---|---|
| Number of participants | 73 | 52 | | |
| Sex, num of females (%) | 206(56.13%) | 19(36.54%) | 0.01[b] | $P = 0.92$ |
| GA at birth (weeks) | 37-42 | 23-36 | 9.51[a] | $P = 1.95 \times 10^{-21}$ |
| PMA at scan | 37-44 (weeks) | 37-44 (weeks) | -0.71[a] | $P = 0.47$ |
| Birth weight (kg) | 1.82-4.80 | 0.59-3.16 | 8.70[a] | $P = 3.22 \times 10^{-18}$ |
| Time points | 2300 | 2300 | | |
| Scan time (mins) | 15 | 15 | | |

[a]Z (Mann–Whitney U test).
[b]$X^2$ test.
